# How economic exchange can favour human genetic diversity

**DOI:** 10.1101/2025.08.21.671532

**Authors:** Cedric Perret, Luís Santos-Pinto, Laurent Lehmann

## Abstract

By allowing individuals to use goods they do not produce, economic exchange is recognised as driving the wide diversity of economic activities seen in human societies. Since productivity also depends on innate abilities, we ask whether economic exchange could have influenced human evolution and promoted adaptive genetic diversity. We model a system where individuals produce and exchange goods under a Walrasian equilibrium, with abilities determined by an evolving quantitative genetic trait. We then analyse how exchange shapes the evolutionary pressures on this trait. Our analysis demonstrates that exchange consistently promotes negative frequency-dependent selection, which favours the maintenance of genetic diversity. Exchange also generates stable long-term adaptive polymorphism when the production of goods requires different abilities. Importantly, we establish that the mode of exchange matters: markets, where individuals can switch trading partners, promote genetic diversity under broader conditions than when exchange occurs in isolated pairs. Finally, we show that genetic diversity and economic specialisation can facilitate the emergence of the other under a wider range of conditions. Our findings suggest that economic exchanges play a crucial role in fostering biological diversity and offer insights into how a culturally determined mode of organisation may have shaped human evolution.

## 1 Introduction

The practice of exchanging goods and services is one of the hallmarks of human societies [1, 2] and has profoundly impacted the social organisation of human groups. The development of economic exchange played a central role in state formation [3, 4], influenced subsistence strategies [5], and, perhaps most notably, is regarded as the primary cause of economic specialisation, leading different individuals to specialise in producing distinct goods and services [6, 7]. From its earliest origins, human history has been intertwined with exchange, as evidenced by archaeological findings of long-distance exchange dating back to the emergence of *Homo sapiens* 200,000 to 300,000 years ago [8]. Then, could economic exchange shape other characteristics of a population beyond its social and economic organisation? If exchange has driven individuals to specialise in producing certain goods, it may have also shaped the evolution of biological heritable traits associated with producing these goods. Indeed, a wide range of traits are likely to affect individuals’ capacities to produce resources [9]. For instance, among the Tsimane people, individuals with strong visual acuity and high running speed appear to be more likely to return from hunting trips with meat, making them more effective food producers [10]. Bajau people of Southeast Asia are able to spend considerable time fishing at great depths of up to 70 meters, a capacity enabled by physiological adaptations that likely increase their catch volume over time [11]. More generally, individuals’ earnings and, presumably their productivity correlate with the main dimensions of personality traits [12, 13], which, like physical traits, are partially heritable [14].

These biological traits evolve by natural selection through their effects on survival and reproduction. Since economic exchange alters the benefit of producing certain goods, it can also affect the reproduction and survival of individuals producing these goods, and therefore change the selective pressures on the traits involved in production. For instance, in the same way that exchange drives diversity in economic activities, are there conditions under which exchange would lead to the emergence of adaptive diversity in genetic traits? This possibility cannot be taken for granted. Evolutionary changes in genetic traits are governed by particular dynamics, with changes generally occurring gradually over multiple generations, in contrast to economic processes, where individuals can freely and nearly instantly adopt strategies when it benefits them. As such, genetic diversity is thought to be favoured only in restricted cases where parents and offspring occupy the same economic roles across generations [15], while arguments proposing that exchange and specialisation lead to phenotypic diversity typically apply to plastic behavioural traits or skills acquired through learning [16, 17, 18, 19]. Yet debate persists [20], and it remains an open question whether adaptive genetic diversity—and, by extension, phenotypic diversity across a wider range of traits—can be favoured by economic exchange in diverse societies, or whether it requires stricter conditions, such as individuals exclusively dedicating their time to distinct roles.

It also remains unclear whether biological traits, such as physical aptitudes or personality differences, influence the conditions under which individuals tend to specialise in producing distinct goods in the presence of economic exchange. The importance of these interactions—between biological traits and humanly devised modes of organisation—is increasingly recognized, as shown by the development of the gene-culture co-evolution theory [21, 22] and the recent integration of genetic data to explain economic outcomes [13, 23, 24]. Examining the interaction between genetic diversity and economic specialisation has also been prompted by recent evidence showing that the emergence of distinct economic roles happened earlier in areas where migration events have led to greater neutral genetic diversity [4, 25].

Given these open and complex questions, a theoretical analysis appears relevant and even necessary to understand whether economic exchange might influence adaptive genetic diversity and whether this changes the relationship between exchange and economic specialisation. Yet, previous studies have tended to stay within the domains of their specific disciplines and did not explore the interaction between exchange, adaptive genetic diversity, and economic specialisation. For example, models focusing on the evolution of genetic diversity under various modes of social organisation have not explicitly considered the role of exchange [26, 27, 28, 29]. Conversely, the Ricardian model of economic specialisation assumes that the abilities of individuals to produce one good or another are exogenous and constant parameters [30].

We address these open questions by building a comprehensive mathematical model that combines standard economic and evolutionary approaches. We examine a population of individuals who produce and exchange goods, explicitly modelling exchange within the Walrasian general equilibrium framework [30, 31]. In addition, the quantities of goods individuals produce depend on (a) their innate abilities, modelled as a quantitative trait and (b) their allocation of time to produce each good, modelled as a behavioural strategy. We use this model to investigate (i) how much exchange can affect the adaptive dynamics of genetic traits and (ii) how genetic diversity and economic specialisation interact. We do this under two scenarios that reflect distinct modes of exchange experienced by humans at different periods in history: individuals exchanging goods in isolated pairs, and individuals engaging in exchange within a large market. We also compare both scenarios to a baseline case of autarky.

## 2 Model

### 2.1 Biological scenario

We consider a randomly mixing population of constant and large size *N* where individuals have economic interactions according to the following non-overlapping generation life cycle. (i) Each individual produces up to two goods, one of type × and the other of type y. (ii) Individuals can exchange goods with each other. (iii) Individuals produce a large number of offspring proportional to their payoff, which depends on the amount of goods they consume after exchange. (iv) Adults die and offspring compete for the breeding spots vacated by the adults and mature, i.e. Wright-Fisher population process [32].

We next describe each of these events assuming asexual reproduction, which simplifies the notation and clarifies the exposition. However, the model and results also apply to sexual reproduction (detailed in Section 3.4), in which case stage (iii) of the life cycle involves random mating among individuals.

#### Production

The quantity of each good that individuals produce depends on (i) an evolving genetic quantitative trait affecting their innate abilities to produce the good and (ii) the time they allocate to the production of the good. Formally, an individual *i* with genetically encoded trait *τ*_*i*_ ∈ ℝ and time *h*_*i*_ ∈ [0, 1] allocated to produce good x, produces an amount *q*^x^(*τ*_*i*_, *h*_*i*_) and *q*^y^(*τ*_*i*_, *h*_*i*_) of goods × and y, respectively. These quantities are assumed to take the following functional forms:

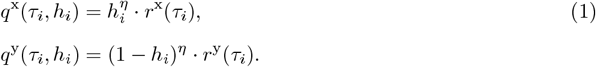

The first component in the products on the right-hand sides of eq. (1) describes the returns of allocating time to produce each good. The parameter *η* ∈ (0, 1), which we refer to as the elasticity of scale, quantifies the decreasing returns to scale of allocating more time to the production of each good. A low value of *η* describes production with strong decreasing returns to scale, while *η* close to one represents constant returns to scale.

The second component on the right-hand sides of eq. (1), the quantity *r*^*k*^(*τ*_*i*_), captures the dependence of the production of good *k* ∈ *{*x, y*}* by individual *i* on trait *τ*_*i*_ and is given by

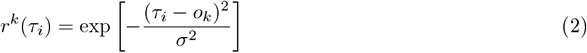

(see top panel of Figure S6 for an illustration). The parameter *o*_*k*_ represents the optimal trait value for which the production of good *k* is maximal. Different values of *o*_*k*_ for different goods capture the notion that these goods require distinct abilities and thus trait values. For instance, high muscle mass may benefit tasks requiring strength such as throwing or sprinting–potentially useful in hunting–but can increase fatigue in endurance activities, thereby limiting the ability to swim or dive for extended periods. The parameter *σ*^2^ *>* 0 represents the production breadth of goods, that is, how deviations from the optimal trait value decrease production. A low *σ*^2^ describes a knife-edge production breadth where a small deviation from the optimal trait causes a sharp drop in production, meaning that production requires highly tailored abilities—for instance, some prey may be inaccessible without the exact strength-endurance balance needed to dive to certain depths. A high *σ*^2^ represents a plateau production breadth where a large deviation from the optimal trait value causes a modest drop in production and so individuals with a wide range of traits can effectively produce each good.

#### Exchange

Individuals can exchange the goods they produce with others by giving up a quantity of one good in return for another. Individual *i* producing quantity *q*^*k*^(*τ*_*i*_, *h*_*i*_) of good *k* and exchanging 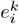 units of this good ends up consuming 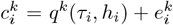 units of good *k*. The quantity exchanged 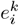 is negative for an individual that gives up some good (“sell” the good), positive for an individual that receives it (“buys” the good), and 0 if no exchange takes place. The quantity of good y given in exchange of one unit of good × is defined as the price *p* of good x.

We consider two possible modes of exchange on top of a baseline situation where individuals live in *autarky* and do not exchange with each other. First, *dyadic exchange* where individuals are randomly paired and each pair is isolated from all others, meaning that they each settle on a potentially different price. This scenario describes early human societies where the large distances between human groups would have limited the number of partners with whom to exchange [33]. Second, we consider *market exchange*, where individuals can freely switch trading partners. This scenario reflects later societies, where exchanges occurred in large markets, as it has been described in many different preindustrial societies [2, 34].

#### Payoff

After exchange, the payoff *π*_*i*_ of individual *i* depends on the amounts of the two goods he consumes according to a Cobb–Douglas form: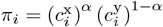. Hence, the goods are complementary and both need to be consumed to entail a positive payoff. The parameter *α* ∈ (0, 1) measures the percentage change of payoff from a change in consumption of good x, holding constant the consumption of good y, and can thus be thought of as a measure of the importance of good × for payoff.

Our aim is to conduct an evolutionary analysis of this model under autarky, dyadic exchange, and market exchange. To do so, we first need to determine the outcomes of the economic interactions that determine individuals’ payoffs. Because the evolutionary dynamics unfold more slowly than individuals’ behaviour within a generation, we assume that prices and quantities settle at an equilibrium when determining payoffs.

### 2.2 Behavioural equilibrium

We assume that individuals are producer-consumers that maximise their payoffs over three decision variables: the quantities consumed of the two goods and the time allocated to produce each good. Thus, the decision problem that consumer-producer *i* with trait *τ*_*i*_ and time allocation *h*_*i*_ faces is

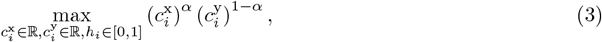

subject to the budget constraint

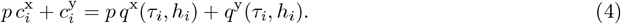

The left-hand side (LHS) of eq. (4) describes the value of the goods individual *i* consumes measured in units of good y (the price of good × is *p* and the price of good y is normalised to 1), while the right-hand side (RHS) represents the value of the goods individual *i* produces. Empirical examples of individuals acting as if they maximise a utility function over quantities of goods are well documented, from experiments in which participants choose among combinations of goods to diet-choice studies (see Appendix A.1).

Solving the optimization problem (3)-(4) provides the equilibrium quantities of each good, 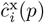 and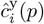, and the time allocation *ĥ*_*i*_(*p)* that maximise the payoff to individual *i* for a given price *p*. Note that this procedure determines the quantity 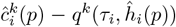 of good *k* the individual is willing to exchange at this price. To close the model, we thus need to determine the equilibrium price. We assume that the price at equilibrium 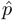 is the one at which the quantities of a good that interacting individuals want to sell (or buy) are exactly equal to the quantities that other individuals want to buy (or sell). In other words, we assume a Walrasian equilibrium—the fundamental equilibrium concept of exchange economics—whereby no surplus nor shortage of resources occurs [35, 30]. Beyond this assumption, we remain agnostic about the mechanism by which the equilibrium is obtained. Walrasian equilibrium has been shown to arise theoretically under different bargaining procedures in dyadic exchange [36, 37], in a wide range of conditions in markets [30] and when price is a culturally evolving trait [38, 39] (a case where the timescale separation we assume remains justified). Empirically, it has been repeatedly shown to emerge rapidly and reliably in a wide range of settings and across decades of experimental research [40, 41, 42, 43].

At a Walrasian equilibrium, the equilibrium price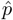, has to satisfy

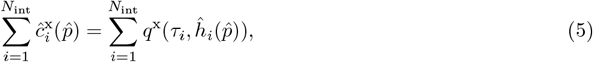

where *N*_int_ is the number of interacting individuals. Under dyadic exchange, *N*_int_ = 2, while for market exchange *N*_int_ = *N*. The solution to the optimisation problem (3)-(4) and eq. (5) determines the equilibrium allocation 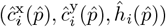 for each individual *i* in the population. At the Walrasian equilibrium, we have the unique feature that all individuals end up consuming precisely the quantities that would maximise their payoff, and thus, we take this consumption to determine their realised material payoff. In turn, and as is standard in evolutionary biology (e.g. 44), we assume that individuals’ reproductive success are proportional to their payoffs.

We solve the behavioural equilibrium explicitly for the two modes of exchange along the special case of autarky in Appendix A to obtain the quantities consumed for each good, the time allocation and the price at equilibrium for any distribution of types of individuals. Having established the equilibrium behaviour as a function of individual traits, we can now proceed to the evolutionary analysis.

### 2.3 Evolutionary analysis

We make the standard assumptions of quantitative trait evolution that the mutation rate is low and the effect size of mutations on trait values are small (e.g. 45, 46, 47). With these assumptions, we can characterise long-term gradual evolution by focusing on the payoff *π*(*τ, θ*) of a single individual with trait *τ* —henceforth called a mutant—which arises in a population of individuals that are monomorphic for trait *θ*—henceforth called the resident population (method detailed in Appendix B.1). According to our assumptions, the payoff of a mutant, derived in Appendix B.2, is

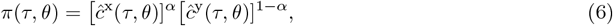

with equilibrium consumption of goods × and y given by

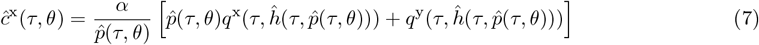

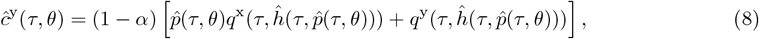

where 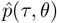 is the equilibrium price a mutant individual faces, which satisfies

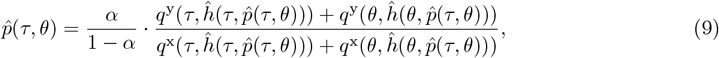

under dyadic exchange and

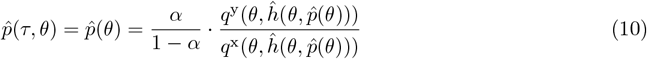

under market exchange. Finally,

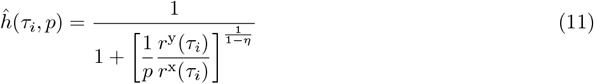

denotes the equilibrium time an individual with trait *τ*_*i*_ ∈ *{τ, θ}* facing price *p* allocates to produce good x.

At the behavioural equilibrium, the quantities consumed, *ĉ*^x^(*τ, θ*) and *ĉ*^y^(*τ, θ*), consist of fractions of the total budget an individual can consume measured in units of good y (expression in square brackets in eqs. 7–8) and translated into quantities of goods consumed via prices. These fractions are the importances for payoff of each good, *α* for good × and (1 − *α*) for good y. The time 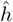 devoted to producing good × at equilibrium increases with the price of good *x* (noting that the price of good y is normalized to 1) and with the individual’s relative efficiency in producing × versus y. The strength of this relationship depends on *η*: when *η* is close to 1, time allocation is highly responsive to changes in price and productivity, whereas as *η* approaches 0, time allocation becomes nearly independent of both price and individual traits. Finally, the price at the Walrasian equilibrium depends on (i) how much good × contributes to the payoff relatively to good y, and (ii) how rare good × is relative to good y—between the two individuals in dyadic exchange or among the resident population in exchange, where the mutant is ignored due to large group size.

We now infer the evolutionary dynamics from eqs. (6)–(10) as follows (see Appendix B.1 for more details). First, the population will evolve by directional selection with the average trait value in the population changing in the direction indicated by the payoff gradient, *∂π*(*τ, θ*)*/∂τ* |_*τ*=*θ*_, until a singular point *θ*^*^ is reached where the payoff gradient vanishes. Once the population has converged this trait value *θ*^*^ under directional selection, i.e. a so-called “convergence stable” trait value, at which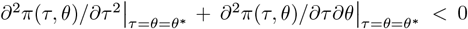 must hold, the nature of selection at that point can be of two types. Either (a) selection is stabilising so that the trait value is locally uninvadable and evolution stops; or (b) selection is disruptive, in which case evolutionary branching occurs and the population evolves further. Disruptive selection requires that the singular point is a payoff minimum, 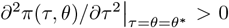, which entails that the population will subsequently divide into two distinct morphs with different trait values (polymorphism), i.e. evolutionary branching occurs. It follows from these considerations that polymorphism evolves by gradual evolution if directional selection is first convergent and subsequently disruptive. A necessary condition for this to occur is thus that the mutant payoff increases when mutant and population trait values move in opposite directions at a singular point: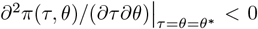. This condition is further necessary and sufficient for two types that are symmetrically diverged around a singular point to mutually invade each other and thus coexist in a protected polymorphism by negative frequency-dependent selection [45].

We divide our evolutionary analysis into two parts to distinguish between the role of exchange and time allocation in favouring genetic diversity. First, we analyse the effect of exchange on adaptive genetic diversity holding time allocation fixed and set *q*^*k*^(*τ, h*) = *q*^*k*^(*τ*). Second, we let the time allocation depend on the trait (as per eq. 11) and analyse the full model. For each case, we start by analysing the trait value that would be favoured by evolution in the absence of exchange (autarky).

## 3 Evolutionary analysis with fixed time allocation

### 3.1 Autarky

In the absence of exchange, the quantities that an individual consumes are equal to what it produces whereby the mutant payoff in eq. (6) simplifies to

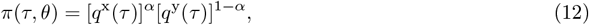

which does not depend on the resident trait value. We show in Appendix B.3.1 that any singular point *θ*^*^ satisfies

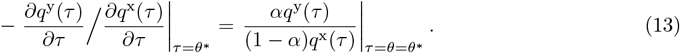

This can be interpreted as saying that the rate at which the production of good × can be substituted for good y (the LHS) is equal to the net relative importances of both goods for payoff (the RHS). We assume for the rest of the analysis of fixed time allocation that the production functions are such that there exist a unique singular point. Since the RHS is positive, this requires the LHS to be positive as well, which in turn implies that the partial derivatives of the two production functions have opposite signs. In other words, we assume that there is a trade-off between the production of the two goods. We also assume the singular point is a maximum of payoff, which altogether entails that under autarky the singular point is a generalist trait that is both convergence stable and uninvadable (see Appendix B.3.1).

Using eqs. (1)–(2) the singular trait becomes

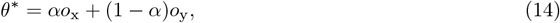

which is located between the two optimal production values. In the absence of exchange, the population thus evolves towards a generalist trait value, leading individuals to produce each good, albeit with reduced efficiency compared to the maximum achievable production for each.

### 3.2 Dyadic exchange

With exchange under dyadic interactions, we find by using eqs. (6)–(9) that the payoff gradient is exactly the same as under autarky, meaning the singular point will also satisfy eq. (13) and be convergence stable. An intuition for this result stems from observing that by substituting eq. (8) into eq. (6), the payoff to a mutant is proportional to (i) the net value of the goods the individual produces, which is affected by its genetic trait through direct effects on produced quantities, and (ii) a factor depending on the price, which is indirectly affected by the genetic trait. It turns out that selective effects stemming from trait effects on the price vanish (eq. A-54), whereby the selection pressure on the trait depends only on direct effects on quantities produced, weighted by the price in a monomorphic population where no exchange occurs. This price (eq. (10)) at the singular point is then equal to the RHS of eq. (13), and so eq. (13) applies to exchange.

Hence, exchange does not alter the nature of directional selection and the population first evolves towards individuals expressing the same generalist trait as under autarky. Yet, exchange affects the evolutionary stability of the convergence stable trait value (see Appendix B.3.2). To understand this, we first note that the presence of exchange always entails that an individual’s trait increases payoff if it moves in oppositive direction than population trait values:

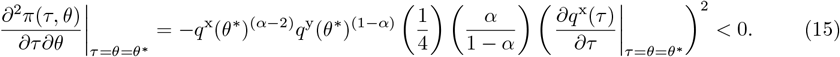

The rationale behind this result is that when individuals from the population deviate from the singular point and thus produce more of one good, that good becomes more common, leading to a decrease in its price, while the other good becomes rarer and more valuable. As a result, an individual whose trait deviates in the opposite direction produces more of the now scarcer good, and achieve a higher payoff than those who continue producing the more abundant good. This establishes that economic exchange creates conditions favouring the maintenance of different traits by selection (see also Appendix B.3.2), but it does not ensure that selection will favour distinct morphs from a monomorphic population under exchange. This second statement is true if selection is disruptive at *θ*^*^, as it means that mutants deviating from the mean have a higher payoff than those at the mean. We find that disruptive selection occurs when

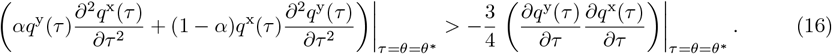

This requires that the second derivatives of the quantities produced with respect to the trait tend to be positive and large enough (at least one derivative must be positive as the RHS is positive, since the partial derivatives are of opposite signs at the singular point, see eq. 13). Disruptive selection thus requires that deviations from the singular point lead to sufficiently high increasing returns to scale in good production (the production function is convex in trait values around the singular point).

The results so far are generic, applying to any production function and we now turn to fully work out the analysis by using our explicit production function (eqs. (1)–(2)). The above results imply that the (convergence stable) singular point will be given by eq. (14) and substituting this into eq. (16) gives the condition for disruptive selection

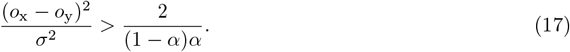

The LHS captures the squared difference between the trait optima *o*_x_ and *o*_y_, scaled by production breadth *σ*^2^. This measures how much an individual excelling at producing one good performs poorly at producing the other and represents the degree of trade-off between producing the two goods. The population becomes polymorphic when this trade-off is strong, as it would be the case for goods that require very different traits to be produced. The RHS of eq. (17) depends on the relative value for fitness of good × compared to good y. Since the denominator is largest when *α* = 0.5, the condition is easier to satisfy when the two goods are equally important for payoff.

We validate these analytical results using individual-based simulations (see Appendix C for details). Simulations illustrate the typical trajectory of trait evolution in a scenario where the population initially lives in autarky and exchange is introduced after 5000 generations. Individuals initially have trait values near the optimum for producing good y, with both goods equally valued for payoff (*α* = 0.5). Before exchange, the population converges on the singular trait value (Figure 1, top left), as predicted analytically. During this time, (i) the population shifts from producing predominantly good y (indicated by a large proportion of red in the bottom left panel of Figure 1) to producing both goods (a combination of blue and red), and (ii) the total quantities of goods produced decreases drastically. This is because both goods are equally valuable for fitness, and in absence of exchange, individuals must produce sufficient quantities of both goods. When exchange is introduced, the population quickly divides into two morphs, with selection pushing the traits of each morph away from each other until each of the morphs has a trait close to the value optimal for production of one of the two goods (top left panel of Figure 1). During this process of diversification, the quantities produced, exchanged, and consumed largely increase, demonstrating the benefits of exchange (bottom left, top right, and bottom right panels of Figure 1). At the long-term evolutionary equilibrium, one fraction of the population produces predominantly good x, while the other fraction produces almost exclusively good y, yet all individuals consume both goods. Exchange allows for genetic diversity, raising production of both goods while maintaining an optimal allocation of goods. This is why the population composed of the two morphs who exchange with each other has an average payoff 13 times higher than a population of generalists who do not exchange (generation 12401 vs. 4801).

**Figure 1:**
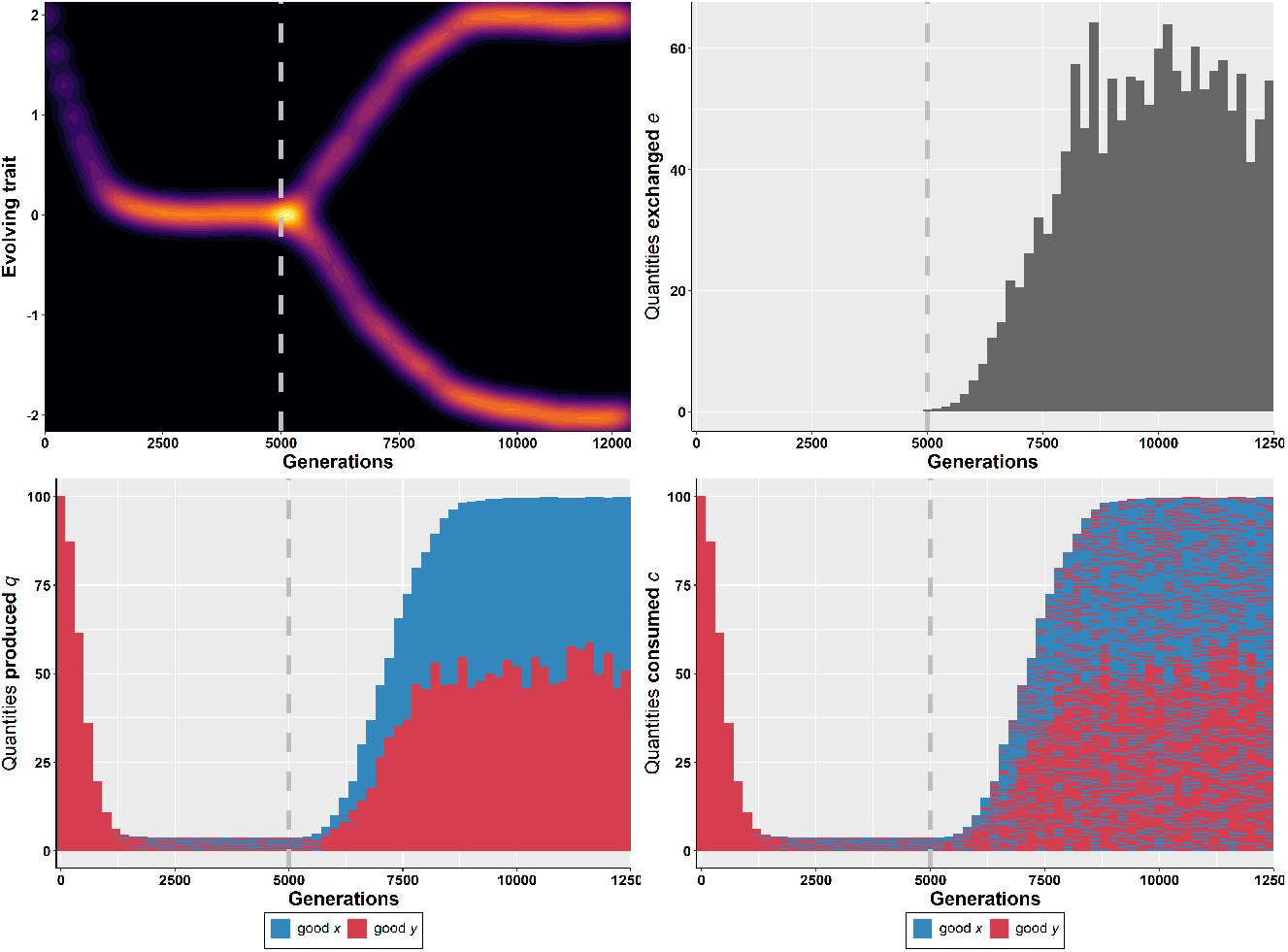
Distribution of trait values (top left), of quantities produced *q* (bottom left), exchanged *e* (top right), and consumed *c* (bottom right) over generations. The population remains in autarky for the first 5000 generations, after which dyadic exchange is introduced (indicated by the grey dotted line). For clearer visualisation, the quantities shown are of a subset of 100 randomly selected individuals, and they are stacked vertically: good × of individual 1 at the bottom, followed by their good y, then good × of individual 2, and so on. Parameters: production breadth *σ*^2^ = 1, equal value of goods *α* = 0.5, optima of production *o*_x_ = −2, *o*_y_ = 2, population size *N* = 5000, mutation rate *µ*_m_ = 0.01 and standard deviation (SD) of mutations *σ*_m_ = 0.02.

Supplementary Figure S7 illustrates the dynamics of the trait when one good is more important for the payoff than the other. It confirms that (i) a population in autarky converges and remain towards a singular point, which is closer to the optimal of production of the most important good, and (ii) diverges once exchange is introduced. At the long-term evolutionary equilibrium, the majority of individuals have a trait close to the optimum of the good that is most valuable for fitness.

### 3.3 Market exchange

We have shown that dyadic exchange (a) consistently promotes, at least temporarily, the coexistence of different morphs and (b) generates stable adaptive genetic diversity when the abilities necessary to produce the goods are different enough. Do these results hold when exchanges take place in a market rather than through isolated dyadic interactions? Under market exchange, the mutant payoff (eq. 6) and the quantities consumed (eq. 8) remain the same as under pairwise interactions, only the price needs to be changed (eq. (10)). Our analysis under market exchange (detailed in Appendix B.3.3) reveals the same qualitative results than under dyadic exchange: (i) the convergence stable singular point is the same as under autarky (given by eq. (13)), (ii) an individual’s trait increases payoff if it moves in oppositive direction than population trait values (see eq. A-67), and (iii) the condition for disruptive selection is satisfied when trait effects result in increasing returns to scale in good production (the production functions are sufficiently convex in trait values) but is weaker and holds under a broader parameter range than under dyadic exchange (see eq. A-69).

Using the explicit production function (eqs. (1)–(2)), the above generic results then imply that under market exchange the convergence stable singular point is again given by eq. (14), but the condition for polymorphism to emerge (derived in Appendix B.3.3) is

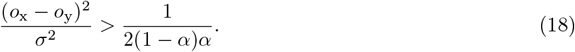

Comparing the RHS of eq. (17) and eq. (18) (and simulations in top panel of Figure S9) illustrate that polymorphism emerges under broader conditions in market than dyadic exchange. This suggests that the development of markets can favour genetic diversity in contexts where dyadic exchange alone is insufficient. The reason for this difference is that in dyadic exchange, genetic diversity is hindered because an individual producing more of a good lowers its price at the same time. As a result, genetic diversity only evolves when the increase in production associated is sufficiently large. However, this effect does not take place in a market, as an individual’s trait does not affect the price.

### 3.4 Sensitivity to model assumptions

We now assess the robustness of our main results to changes in model assumptions. First, we confirm that genetic diversity emerges in the conditions predicted by eqs. (17) and (18), even when mutation effects are larger (Supplementary Figure S8). Second, while we conducted the previous derivations and simulations assuming asexual reproduction for simplicity, our results apply under sexual reproduction (see A-32 for why the analytical results hold generally). We confirmed this by analysing simulations with sexual reproduction where the trait *τ*_*i*_ of individual *i* is the additive combination of the genetics effects of two alleles it carries, assuming random mating (Appendix C). Both dyadic and market exchange continue to generate and maintain adaptive genetic diversity, with the genetic variance in the population increasing over time from the branching point onwards until reaching an equilibrium (Figure S2). The genetic variance is smaller with sexual reproduction as offspring of specialised individuals reintroduce unfit generalist individuals (Figure S1). This should generate selective pressures against the generation or expression of these intermediate phenotypes [48], e.g. assortative mating, but exploring these possibilities lies beyond the scope of this paper.

Finally, while we initially assumed a single allocation trait simultaneously affecting the production of both goods (as usually done in biology [49]), we also analysed a version of the model in which the production of each good depends on a distinct trait, subject to a limit on the resources an individual can allocate to these traits (see Appendix D for models comparison). Once again, under both dyadic and market exchange, exchange promotes genetic diversity. This shows that our results of the emergence of genetic diversity under exchange also applies when distinct traits independently govern the production of each good, provided the reasonable assumption that individuals face a constraint limiting total trait investment.

## 4 Evolutionary analysis with time allocation decision

Thus far, we have examined the influence of exchange exclusively on the evolution of genetic diversity by keeping the time allocation *h* constant. We now examine how exchange shapes both genetic diversity and economic specialisation by letting individuals express their optimal time allocation to producing each good (eq. 11), which itself depends on the evolving trait. We first carry out the evolutionary analysis along the same line as in the previous sections, before examining more closely the interplay between genetic diversity and economic specialisation using simulations. Due to the complex feedbacks introduced by time allocation decisions, we carry out the forthcoming analysis using only the specific production functions defined in eqs. (1)–(2). Furthermore, while some analytical results can be obtained under market exchange, the analysis for dyadic exchange must be conducted numerically, as the equilibrium price cannot be solved analytically. We therefore explore dyadic exchange by varying two parameters: the elasticity of scale *η* in increments of 0.05 and the production breadth *σ*^2^ over the range [1, 20] in increments of 1, while setting *o*_x_ = −2, *o*_y_ = 2 and *α* = 0.5.

### 4.1 Autarky

In autarky, we find that the time allocation at equilibrium entails that all individuals allocate a fraction 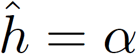of their time to produce good × (Appendix A.2). Substituting this time allocation into the mutant payoff (eq. (12)) and computing the payoff gradient (eq. (A-32)) yields the same single equilibrium point *θ*^*^ = *αo*_x_ + (1 − *α*)*o*_y_ than when time allocation was fixed. Furthermore, we find that this singular point is convergent stable and uninvadable (Appendix B.4.1). In autarky, individuals thus evolve to have a similar genetic trait between the two optimals of production, as found under fixed time allocation.

### 4.2 Exchange

We now conduct the evolutionary analysis in the presence of exchange using eqs. (6)–(10). In the presence of exchange, whether dyadic or market based, we find that there is a unique singular point taking the usual form of eq. (14) (Mathematica notebook for dyadic exchange and Appendix B.4.3 for market exchange). Furthermore, this trait value is convergence stable on the range of *η* and *σ*^2^ values specified at the beginning of section 4. As under fixed time allocation, the population thus first evolves by directional selection towards having the same generalist value of the trait.

At this value, the interaction coefficient is negative on the range of *η* and *σ*^2^ considered for dyadic exchange and is always negative for market exchange (eq. (A-67)), which favours the coexistence of individuals with different trait values. We next examine the conditions under which disruptive selection occurs and thus polymorphism emerges by gradual evolution. For dyadic exchange, we assess these conditions numerically across the parameter space considered and the results are displayed in the left panel of Figure 2. For market exchange, we obtain the following condition (Appendix B.4.3)

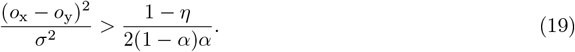

**Figure 2:**
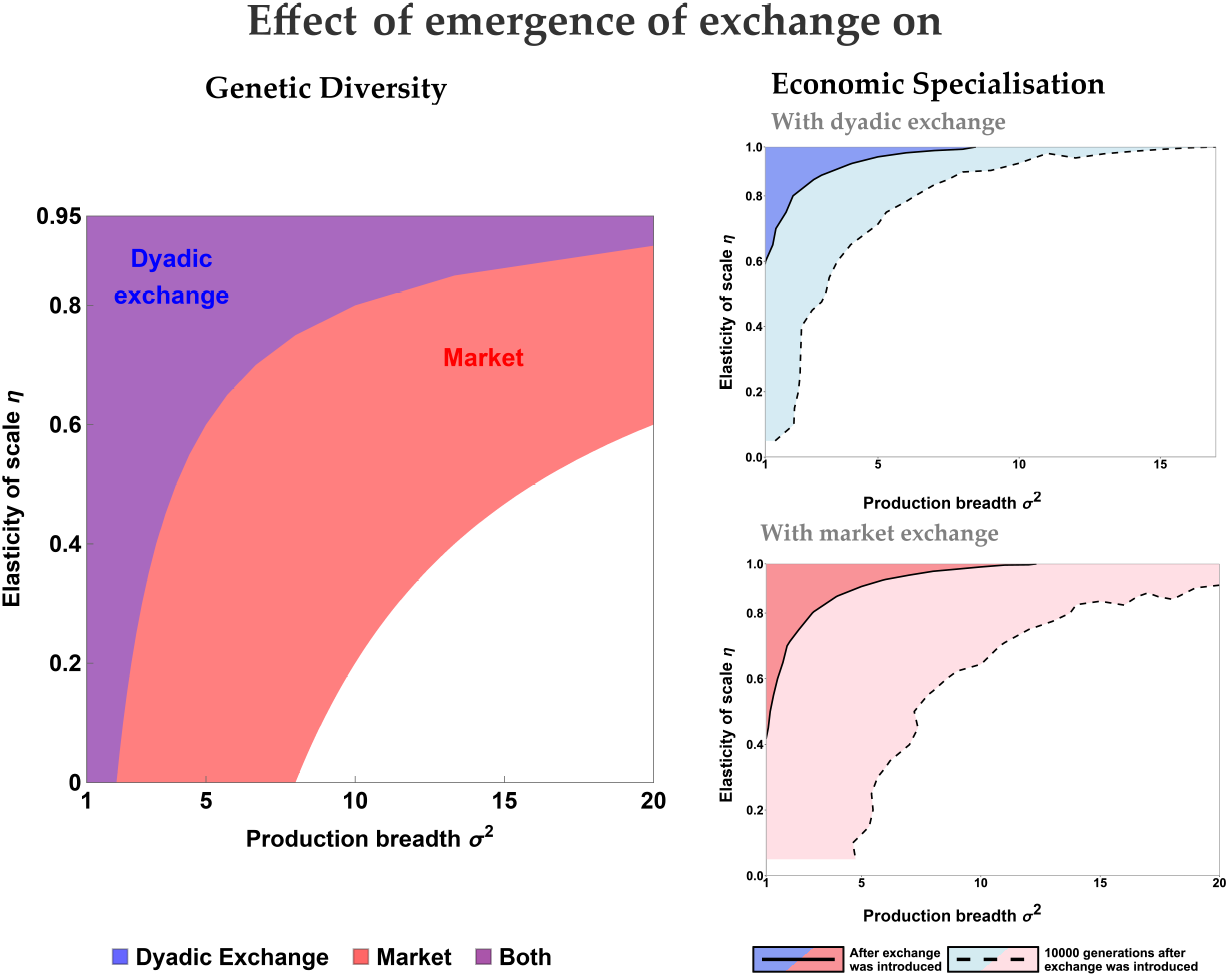
Effect of introducing exchange on genetic diversity and economic specialisation across production breadth *σ*^2^ and elasticity of scale *η*. The left panel shows the conditions under which genetic diversity is predicted to emerge, defined by singular trait values that are both convergence stable and invadable. The right panel shows the conditions under which economic specialisation is observed after 1 (dark shading) or 10,000 generations (light shading) following the introduction of dyadic (top) or market exchange (bottom). Economic specialisation is measured by the average difference between the highest and lowest values of *h* exceeding 0.8 across 10 simulations. Parameters: optima of production *o*_x_ = −2, *o*_y_ = 2, equal value of goods *α* = 0.5, population size *N* = 1000, mutation parameters *µ*_m_ = 0.01, *σ*_m_ = 0.02. Initial population is drawn from a Normal distribution centered on 0 with SD 0.05.

This expression is identical to the one derived under fixed time allocation in eq. (18), except that the RHS now depends on 1 − *η*.

These results show that for both dyadic and market exchange, when *η* is close to 0, the time allocation is largely independent of the evolving traits and selection is disruptive under similar conditions than the ones derived in eqs. (17) and (18), consistent with the results obtained under fixed time allocation. When *η* is higher, meaning when the effect of diminishing returns is less pronounced, the range of values of production breadth *σ*^2^ within which polymorphism emerges broadens. When *η* is close to 1, individuals tend to allocate a significant proportion of their time to producing either one good or the other (see eq. (11)). As a result, polymorphism emerges almost inevitably, even when *σ*^2^ is high, that is, even when the production of goods does not require tailored abilities. The key result is that a context favourable to economic specialisation significantly broadens the conditions for genetic diversity to emerge. Comparing the conditions for polymorphism to emerge between dyadic and market exchange (Figures 2 and S9) reveals that polymorphism arises under a broader range of conditions in market exchange than in dyadic exchange.

### 4.3 Interplay between economic specialisation and genetic diversity

We now finally examine how the interaction between exchange and genetic diversity affects the distribution of time allocation in the population. To do so, we start by illustrating the evolutionary dynamics using simulations, again introducing exchange after a period of autarky (Figures S10–S11). During the period of autarky, the traits evolve towards a generalist value and individuals divide their time equally between producing the two goods (fraction *h* = *α* = 1*/*2 of their time on × and y). The introduction of exchange then leads individuals to shift their time allocation from producing both goods to focusing more on a single good. In other words, individuals begin to economically specialise. As generations proceed, two morphs emerge and diverge genetically, and along this process, individuals increasingly allocate their time to the good for which their genetic trait is better suited (as predicted by eq. (11)). At equilibrium, the population consists of individuals with distinct genetic traits and distinct time allocation patterns (Figures S10–S11).

How in this process does genetic diversity and economic specialisation reinforce each other? To better understand this, we examine economic specialisation when exchange is just introduced, and then after 10,000 generations. First, upon the introduction of exchange and as the population is still genetically homogeneous, the extent to which economic specialisation emerges depends strongly on parameter values. It ranges from full specialisation when returns to scale are high and production breadth is low, to individuals continuing to divide their time equally between goods when the opposite is true. Importantly, the parameter combinations that favour economic specialisation (in particular the range of *η* values) are also those where genetic diversity consistently emerges (Figure 2), indicating that economic specialisation promotes genetic diversity, as suggested analytically by eq. (19). Yet full economic specialisation, in the sense of individuals engaging in entirely distinct production activities is not required for genetic polymorphism to emerge.

Second, our results show that after evolution has unfolded and genetic diversity has potentially emerged, economic specialisation is observed across a broader range of parameter combinations than immediately after the introduction of exchange (Figures S12–S13). Notably, the conditions under which economic specialisation is ultimately observed closely overlap with those that initially favoured the emergence of genetic diversity (Figure 2). This illustrates that the emergence of genetically diverse morphs can in turn promote economic specialisation (as shown by eq. (11) and displayed in Figure S10), and that in some cases, genetic diversity enables the development of economic specialisation that would not have occurred otherwise.

## 5 Discussion

The impact of economic exchange on the organisation of human societies has long been recognised, yet its possible footprint on innate human abilities is less understood. In this paper, we introduced an individual producer-consumer model that we combined with a standard economic model of exchange and a standard evolutionary model to investigate how the presence of economic exchange might shape the selective pressures acting on a quantitative trait involved in the production of goods. In our model, the evolving traits can represent a wide range of biological characteristics, provided they have a heritable genetic component and are subject to a trade-off, either because they influence the abilities to produce multiple goods and/or because they draw on a shared, limited resource budget. This includes most morphological and physiological traits, as well as a substantial proportion of behavioural traits [9, 50]. Our findings are summarised in the following key points.

First, our theoretical analysis reveals that the introduction of exchange can shift the nature of selection pressures, from maintaining by stabilising selection an homogeneous population of generalists under autarky, to favouring adaptive genetic diversity. Specifically, with exchange, selection (i) consistently promotes, at least temporarily, the coexistence of different morphs through negative frequency-dependent selection, and (ii) can generate stable long-term adaptive polymorphism by disruptive selection. This shift occurs because exchange allows individuals whose heritable traits make them efficient at producing a single good, to still access the other good, and thus to have a higher survival and reproduction. At the same time, the price dynamics—where an increase in production of a given good lowers its relative value and, consequently, the fitness of these producers—prevent any single type from completely replacing others. Importantly, these processes, despite resulting in an increase in the average payoff in the population, do not require conditions such as population structure, competition between groups, or groups facing different environments.

Second, we identify the conditions under which exchange promotes adaptive genetic diversity. As we have seen, a polymorphism is favoured by selection when small deviations from a generalist trait lead to disproportionately higher production of one good, meaning the gain in production of one good outweighs the loss in production of the other. This translates in our model in three conditions. First, production of the goods must require sufficiently different innate abilities, that is, the trait values that maximise production of each good, must lie sufficiently far apart. Second, the goods must display a knife-edge production breadth (small enough *σ*^2^) where small deviations from the optimal trait values (*o*_x_ and *o*_y_) lead to large reductions in quantities produced. Third, one good can not be much more valuable than the other in terms of fitness, that is the importance of good × for payoff (*α*) cannot be too close to 0 or 1. Our results further show that when exchange takes place through a market involving many participants, selection generates stable polymorphism in a wider range of conditions. This suggests that the transition from localised, bilateral exchanges between individuals to broader market economies, could have resulted in an increase in human genetic diversity.

Third, our model clarifies how genetic diversity—variation in individuals’ innate productive abilities— and economic specialisation—variation in time individuals allocate producing each good—can jointly change. The interactions between adaptive genetic diversity and economic specialisation have been hard to study as they can feed back into each other, but our theoretical approach allows us to disentangle their roles. The presence of economic specialisation on top of economic exchange greatly widens the range of conditions under which natural selection generates adaptive genetic diversity. This extends the verbal argument that different economic and social roles promote phenotypic diversity through learning [17, 18, 19] (i) to include phenotypic diversity arising from adaptive genetic variation, as proposed by [20], and (ii) to include traits that are more rigidly determined by genetic factors. However, our results also clarify that different economic roles are not always necessary for genetic diversity to emerge, as the presence of economic exchange can result in genetic diversity, even when individuals do not differ in the time they spend producing one good or the other. In contrast, it can even be the emergence of genetic diversity which allows for the development of different economic roles.

Identifying the conditions and processes that favour genetic diversity has long been a central question in evolutionary biology, as it underpins the adaptability of populations to environmental changes and provides insights into past evolutionary processes. Importantly, it is well understood that cultural phenomena could affect genetic diversity, as for instance shifts to patrilineal descent resulted in a large decrease in the Y-chromosome diversity [51]. Yet, such examples remain rare. Our work adds a new example of this relationship by showing that the development of economic exchange, and the factors that allowed it [52], such as changes in technology, infrastructure and population density, could have resulted in significantly increased genetic diversity. So far, a correlation between exchange and genetic diversity has only been tested indirectly, as previous studies have examined correlations between neutral genetic diversity and economic specialisation rather than adaptive diversity and exchange directly [4]. Integrating more detailed data on economic systems could allow for a more direct test. Alternatively, a population such as the Tsimane people presents a particularly interesting case study, as different groups vary in their market integration, and this variation has already been shown to influence the social structure of these groups [53].

In economics, specialisation was first considered a direct and almost inevitable consequence of eco-nomic exchange, as illustrated by the famous quote by Adam Smith: “the division of labour is limited by the extent of the market.” Yet, since Adam Smith, a large body of evidence has shown that this statement should be tempered, as past and current human societies exhibit wide variability in their levels of specialisation. While specialists are prevalent in early states [54], small-scale societies tend to have very limited division of labour, aside from gendered divisions [55]. Therefore, a crucial question is what factors could account for the emergence of specialisation in some societies, while it remains absent in others. Previous hypotheses usually focus on the role of changes in technology, group size, or political elites, and ignore the potential role of biological factors. Recently, [4] have proposed that genetic diversity could have been an important factor, based on evidence that economic specialisation emerged first in groups with higher genetic diversity. Our analysis supports this verbal model and confirms that there are scenarios where economic specialisation could emerge only under conditions favouring genetic diversity first. In addition, our findings show how such diversity can arise through disruptive selection, and identify some of the conditions under which this occurs.

Essentially no model in the literature has previously explored the conditions under which adaptive diversity can emerge in heritable traits related to the production of goods when economic exchange is modelled explicitly, as we have done here. Previous models have demonstrated that diversity in an evolving trait is expected when social interactions favour anti-coordination [26, 27, 28]; for example, tasks where two individuals performing different actions achieve a higher payoff than two individuals performing the same action [26]. Yet, these models focus on how heritable differences can be maintained given a particular fixed payoff structure. In contrast, our model takes a different perspective by identifying exchange itself as the process which generates this payoff structure under which specialisation and genetic diversity emerge. In line with this distinction, our model is particularly related to that of [29], who show that the custom of sharing goods can also promote genetic diversity. However, their results depend on having sufficiently large groups, whereas our results show that genetic diversity could emerge with economic exchange regardless of group size. This raises the broader question of how transitions from food-sharing practices to market-based exchanges may have influenced the evolution of adaptive genetic diversity.

Our model may seem to overlook the fact that individuals’ abilities to produce a given good is also determined by traits that are acquired through learning or experience, rather than inherited biologically. However, even if learned skills contribute significantly to certain productive abilities, as long as skill acquisition is independent of the evolving genetic traits under focus, selection would still act on the heritable component of these traits as described by our results. Exploring more complex interactions between genetic and learned traits, where individuals learn skills that are either poorly or well suited to their genetic traits, could be an interesting extension for future work. Another avenue for future work would be to explore cases where prices deviate from those calculated here, for instance, when they are set by political authorities or when the payoff also integrates social norms or outdated preferences.

When Adam Smith famously proposed that exchange leads to specialisation, he asserted that “the difference between the most dissimilar characters, between a philosopher and a common street porter, for example, seems to arise not so much from nature, as from habit, custom, and education” [6]. Two centuries later, it is widely accepted that biological and environmental factors interact in shaping human traits, making the picture far more complex than Smith initially envisioned. Our study extends this insight by demonstrating that exchange itself—the very mechanism Smith described—not only influences learning and skill acquisition, but may also have shaped adaptive genetic diversity through evolutionary pressures. The historical transitions from simple exchange to complex markets may not have merely reorganised production but actively reinforced biological differentiation, making exchange a long-term evolutionary pressure.

## Acknowledgments

We thank Ingela Alger, Paul Seabright, members of the Institute of Advanced Studies in Toulouse, and of the Department of Economics at the University of Lausanne, for their valuable feedback and insightful comments.

## Appendix

### Appendix A Behavioural equilibrium

In this section, we derive the behavioural equilibrium that determines how individuals allocate time, produce, and exchange goods. We begin by stating the behavioural assumptions and motivating them empirically, and then solve the individual optimisation problem (3)–(4) introduced in the main text.

#### Appendix A.1 Behavioural assumptions

To compute the behavioural equilibrium, we assume that individuals are producer-consumers who choose consumption of two goods and their time allocation to maximise a payoff function defined over the quantities they consume.

This assumption is supported by a large empirical literature documenting behaviour where individuals act as if they maximise a utility function defined over the quantities of different goods. For instance, revealed preference experiments show that people make consistent choices between bundles of goods, thus behaving as if they are maximising a stable utility function based on the quantities of those goods [1, 2]. A similar pattern is observed when examining how individuals allocate their time to producing goods in more naturalistic settings. For instance, [3] showed that when Agta communities transition from foraging to farming, individuals systematically reallocate time toward more subsistence labour and away from leisure. The fact that most individuals adjust in the same direction is consistent with behaviour as if maximising a similar utility across individuals. Finally, a more biologically grounded example comes from studies of human diet. When choosing among foods, people adjust both quantities and mix to balance protein and calories intake [4, 5].

These decisions, whether about bundles of goods or diet composition, translate outside the laboratory into choices over how much to acquire of different goods through production or exchange, as examined in our model. Importantly, these behaviours do not require advanced cognition or awareness of the utility function, as for instance diet choice have even been documented in taxa such as birds [6]. Rather, these behaviours can be thought to arise through trial-and-error learning processes, instead of through explicit computation by the individual, as is typical of human behaviour in laboratory settings [7].

#### Appendix A.2 Autarky

In autarky, the optimization problem (3)–(4) faced by the producer-consumer individual *i* can be reduced to simply choosing the optimal time allocation and is thus defined by

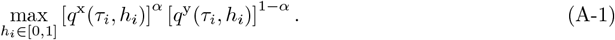

The necessary first-order condition for an interior optimum is obtained by solving the following equation for *ĥ*_*i*_

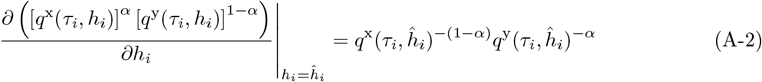

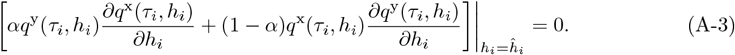

Since the first factor on the right-hand side is strictly positive, the first-order condition reduces to solving the second line for *ĥ*_*i*_.Then, substituting eq. (1) and simplifying gives

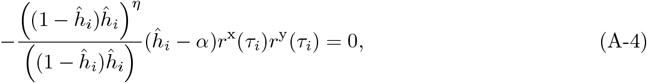

which yields the interior solution

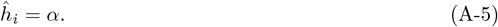

Since the payoff is concave in *ĥ*_*i*_, this is also the optimal solution. Hence, in autarky, each individual allocates a fraction *α* of their time to producing good x.

#### Appendix A.3 With exchange

##### Appendix A.3.1 Producer-consumer decision problem

In the presence of exchange, recall from section 2.1 that the decision problem (3)-(4) that consumer-producer *i* with trait *τ*_*i*_ and time allocation *h*_*i*_ faces is

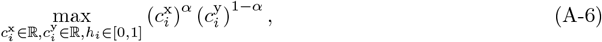

subject to the constraint

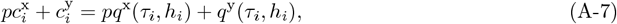

where 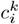 is the quantity of good *k* ∈ *{x, y}* consumed by individual *i, q*^*k*^(*τ*_*i*_, *h*_*i*_) is his “endowment” of good *k* ∈ *{x, y}*, which depends on the time allocation *h*_*i*_, and *p* is the price of good x.

To solve this problem we use standard constrained maximization method [8], and form the associated Lagrangian function

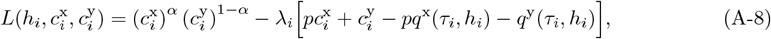

where *λ*_*i*_ is a Lagrange multiplier. A necessary condition to solve the decision problem is that the Lagrangian 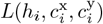 is maximised with respect to the choice variables 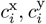 and *h*_*i*_. Using eq. (1), the necessary first-order conditions characterising the solutions to the problem are

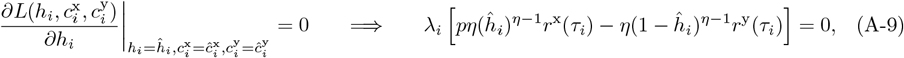

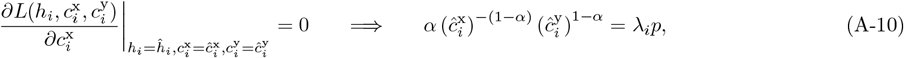

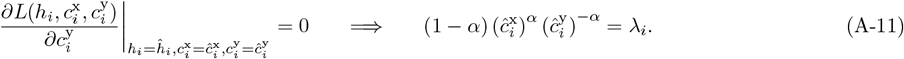

###### Time allocation

To solve the first-order conditions, we first focus on the time allocation. Solving eq. (A-9) for *ĥ*_*i*_ gives which gives

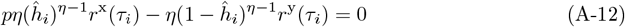

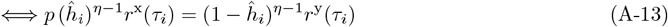

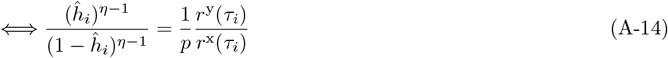

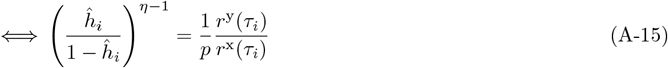

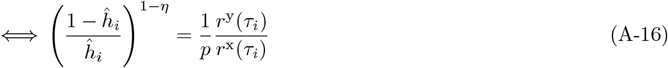

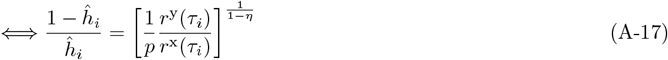

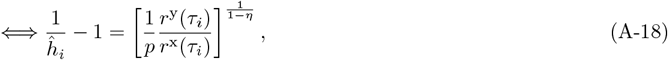

which gives

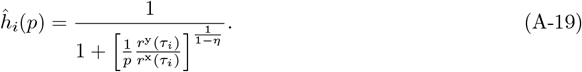

Owing to the fact that 0 ≤ *η <* 1, the Lagrangian is concave in *h*_*i*_, which is sufficient for this solution to maximise the Lagrangian, holding everything else constant. Eq. (A-19) thus describes the time allocation which maximises the payoff of the individual as a function of the price.

###### Quantities consumed

Let us now focus on consumption. Dividing eq. (A-10) by eq. (A-11), we have

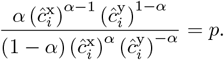

Solving for 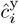 gives

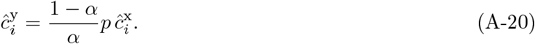

Substituting into eq. (A-7) and solving for 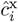, we get the consumption at equilibrium for good × as function of the price

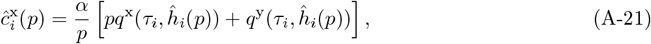

which is called demand function for good × in economics. Substituting back this expression into eq. (A-20), we get the demand function for good y by individual *i*

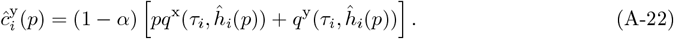

Because the Lagrangian (A-8) is concave in the consumptions 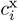 and 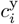and the solutions eqs. (A-21)–(A-22) are unique, these are the optimal solutions to the decision problem [8]. The consumptions so obtained depend on the price, and to close the model we need an expression for the equilibrium price to which we next turn.

##### Appendix A.3.2 Price at equilibrium

Owing to our assumption of a Walrasian equilibrium [9, 10, 11, 12] in section 2.2, the equilibrium price 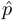 is the one at which the total quantities produced by the interacting individuals equals total consumption.

This requires that the price satisfies

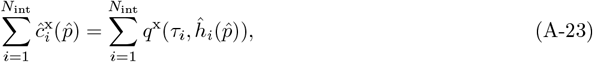

where recall *N*_int_ is the number of interacting individuals, either 2 in dyadic exchange or *N* in market exchange.

Substituting eq. (A-21) into eq. (A-23) and solving for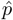, we get the equilibrium price

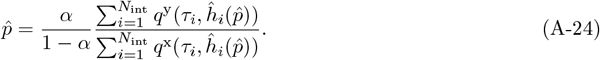

Under market exchange we set *N*_int_ = *N* into this expression. Under dyadic exchange *N*_int_ = 2, there are two interacting individuals, say *i* and *j*, whereby the price can be simplified to

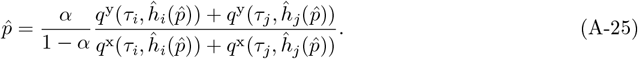

These expressions for the equilibrium price 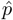 are only implicit, since 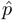 appears on both sides. To solve for the price, we substitute eq. (A-19) along eq. (1) for each individual into eq. (A-24) (or eq. (A-25)) and solve the resulting equation numerically for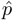. Note that when the time allocation is fixed, the price is directly given by the explicit expressions in eq. (A-24) and eq. (A-25) (since the right-hand sides no longer depend on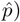. We later show in Appendix B.2.2 that this is also the case under market exchange in a monomorphic population. The above method allows us to determine the equilibrium price, which we next show to be unique.

###### Uniqueness of equilibrium price

We now demonstrate that eq. (A-24) has a unique solution. To do so, we first reformulate eq. (A-24) by defining the function 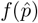 for 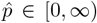 such that solving eq. (A-24) is equivalent to solving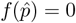. This yields

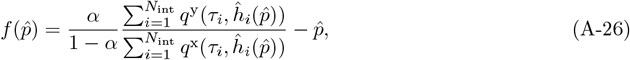

which we show to be a monotonically decreasing function of 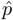 bounded by a positive and negative value.

We first assess the monotonicity of 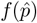. Differentiating 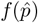 with respect to 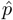 gives

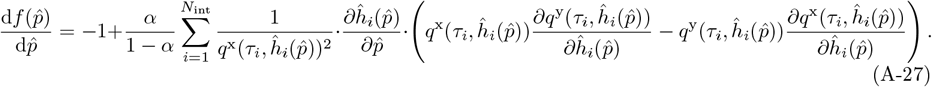

Using eq. (1), the partial derivatives in the brackets satisfy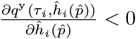, i.e. increasing time allocated to produce good × decreases production of good y, and 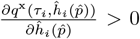, i.e. increasing time allocated to produce good × increases production of good x. Since these signs imply that the bracketed term is negative over the domain of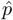, the entire derivative 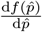 is always negative if and only if 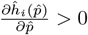 for all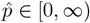. Using eq. (A-19) shows that this is always true as

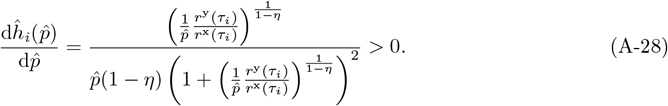

Thus, 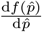 is always negative and the function 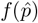 is strictly decreasing.

We next evaluate the sign of the function at the boundaries of its domain. Using the functional forms of 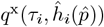 and 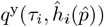 in eq. (1), along 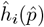 in eq. (A-19), we note that as the equilibrium price approaches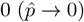, the equilibrium time allocated to produce good × also converges to 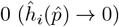, and thus the production of good × vanishes while the production of good y increases to its maximum.

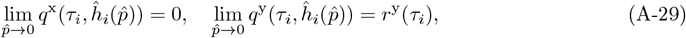

while, as the equilibrium price approaches its maximum 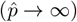, equilibrium time allocated to produce good × converges to its maximum too 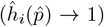, and thus the production of good × increases to its maximum while the production of good y vanishes.

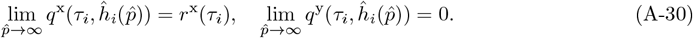

Substituting these limits into 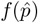 shows that

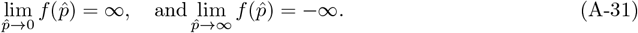

Thus, 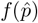 is positive at the lowest bound of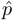, and negative at the highest bound of 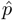. Since 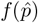 is strictly decreasing, it follows that there exists a unique root, and thus a unique price satisfying *f* 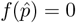.

##### Appendix A.3.3 Equilibrium consumption and time allocation

With the equilibrium price determined, we now turn to the final step, determining time allocation and the quantities ultimately consumed by individuals. The time allocation at equilibrium is obtained by substituting the equilibrium price satisfying eq. (A-24) into eq. (A-19). For consumption, substituting the equilibrium price into eqs. (A-21)–(A-22) gives the demand functions at the Walrasian price. Since at this price, the market clears (eq. (A-23)), there is no surplus or shortage, and all individuals end up exchanging and consuming precisely the quantities specified by their demand functions (this being the unique property of the Walrasian equilibrium). Thus, the quantities consumed at equilibrium for each individual *i* ∈ *{*1, 2, …, *N}* are 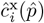 and 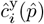, which determine the payoff to each individual. This completes the derivation of the behavioural equilibrium and price, which we can now use in the evolutionary analysis, whether the analytics or the simulations.

### Appendix B Evolutionary analysis

#### Appendix B.1 Adaptive dynamics

To study the evolutionary dynamics of the quantitative trait *τ*, we make the standard assumptions that the mutation rate is low and that the phenotypic effects of mutations are small. Accordingly, we can describe the long-term evolutionary dynamics of the trait by focusing on the invasion fitness (geometric growth ratio) *W* (*τ, θ*) of a mutant with trait *τ* introduced in a population monomorphic for trait *θ* (e.g. 13, 14, 15, 16, 17, 18). We further assume that invasion fitness is of the form *W* (*τ, θ*) = *w*(*π*(*τ, θ*), *R*(*θ*)), where *w* is a monotonically increasing function of payoff *π*(*τ, θ*) and *R*(*θ*) is a one dimensional regulating variable taking density-dependent competition into account and entailing that in a fully monomorphic population the consistency condition *W* (*θ, θ*) = *w*(*π*(*θ, θ*), *R*(*θ*)) = 1 holds for all *θ* ∈ R. Then, it is sufficient to focus on the payoff *π*(*τ, θ*) of a mutant *τ* in a resident *θ* population to characterise the evolutionary dynamics (e.g. 19). The direction in which the trait value changes in a population at *θ* can then be determined from the payoff gradient

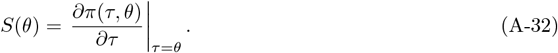

A trait value *θ*^*^ satisfying *S*(*θ*^*^) = 0 will be called a singular trait value and is a candidate evolutionary equilibrium.

The local evolutionary dynamics around singular points can be characterised by the sign of the three components of the following expression:

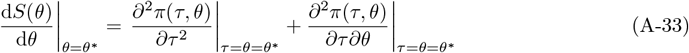

(e.g. 13, 16, 17), which we refer to as the convergence stability, disruptive selection, and interaction coefficients, respectively. The convergence stability coefficient 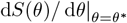 indicates the direction of selection on the trait value when the resident population is in the neighbourhood of the singular point *θ*^*^. A negative coefficient means that a population with a trait higher than the singular point *θ*^*^ will evolve towards a smaller value and thus the singular point is an evolutionary attractor (and vice versa for a lower trait value). The disruptive selection coefficient 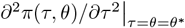 in turn describes what happens when the singular points are reached in the evolutionary process. If this coefficient is negative then selection is stabilising and the population remains at the singular point. If the disruptive selection coefficient is positive then selection is disruptive. If a singular trait *θ*^*^ is both convergence stable and invadable, then it is an evolutionary branching point; namely, an attractor of the evolutionary dynamics that subsequently splits the population into distinct morphs leading to the coexistence of different types in a protected polymorphism (e.g. 16, 17). Branching points are of particular focus in this paper and it is useful to note from eq. (A-33) that a necessary condition for 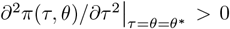 and 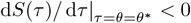 to hold is that the interaction coefficient is negative:

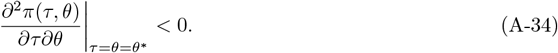

A negative expression implies that individuals achieve higher payoffs when interacting with partners possessing different traits rather than similar ones, thereby favouring the coexistence of different morphs.

The above evolution invasion analysis framework is more general than first meet the eyes for two reason. First, the focus on mutant payoff in a monomorphic resident population to characterise selection does not in fact imply that the evolutionary process needs to unfold in a purely monomorphic population. All the above concepts apply to populations with distributions of traits following standard quantitative genetics modelling assumption, whereby the resident trait *θ* can be always be thought of to be the average trait in a population where variation is segregating through the constant influx of mutations and the trait variance remains small (see 20 for more details on these connections between invasion analysis and quantitative genetics). Second, the focus on asexual reproduction does not in fact imply that the results do not apply to diploid population. On the contrary, assuming that the evolving traits are determined by convex combination of allelic effects that are taken to be the evolving genetic values, the qualitative results of the invasion analysis results also apply to sexual reproduction (see 21 for more details). In summary, all our qualitative results about the characterisation of singular points (their convergence stability, uninvadability, etc…) reported in this paper carry over to sexual reproduction and polymorphic populations, as long as there is no heterozygote advantage in allelic effects. This is indeed confirmed by our simulations (Figures S2, S1 and S5).

#### Appendix B.2 Mutant payoff

We now turn to characterising the mutant payoff *π*(*τ, θ*) required for the invasion analysis under our explicit exchange model. Following our definition in section 2.1, the payoff of a mutant with trait *τ* in a population composed of residents with trait *θ* is

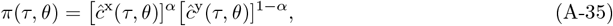

where *ĉ*^x^(*τ, θ*) and *ĉ*^y^(*τ, θ*) denote the mutant consumption of respectively goods × and y at equilibrium. To get these expressions, we adapt the previously derived expressions in Appendix A as follows (where we denote the equilibrium price that the mutant faces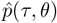, the time allocation of the mutant 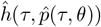 and the time allocation of a resident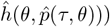.

The equilibrium consumptions of the mutant in eq. (A-35) are obtained by assigning mutant type *τ* to individual *i* in eqs. (A-21) and (A-22) while replacing the price *p* by the equilibrium price faced by the mutant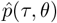 and the time allocation at equilibrium *ĥ*_*i*_(*p*) by the time allocation at equilibrium of the mutant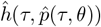. These substitutions yield the expressions in eqs. (7)–(8).

The equilibrium time allocation for mutant and resident for a given price are obtained by respectively assigning type *τ* and type *θ* to individual *i* in eq. (A-19), leading to the expression given in eq. (11).

The equilibrium price under dyadic exchange,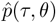 is obtained by assigning type *τ* to individual *i* and type *θ* to individual *j* in eq. (A-25), leading the time allocation 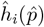 and 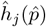 to be rewritten as the time allocation of mutant and resident 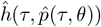 and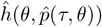. These substitutions yield eq. (9). With market exchange, the mutant *τ* is assumed to have no effect on equilibrium price (since the population is large). The price at equilibrium is then obtained by assigning type *θ* to all individuals in (A-24) with *N*_int_ = *N*, which simplifies to eq. (10) of the main text.

##### Appendix B.2.1 Validity of the monomorphic approximation for the equilibrium price with market exchange

In the case of market exchange, the equilibrium price obtained depends only on the monomorphic population, which may seem counterintuitive since no exchange should take place if all individuals produce the same amount. However, recall that our evolutionary model can be interpreted in a quantitative genetics way such that there is always some variance in trait values in the population owing to the constant influx of mutations (recall the last paragraph of Appendix B.1). The price obtained in the monomorphic population is then an approximation of the price in a population where individuals exhibit variation in trait values. To demonstrate the validity of this approximation, we show that taking variance in trait values in account does not affect the equilibrium price, which still converges to the value calculated under monomorphic assumptions, as long as the population is large enough and the variance in the unimodal trait distribution remains sufficiently small, both of which hold in our model.

Let us now assume, for this section only, that the population trait follows some unimodal distribution with mean trait value *θ* and small variance 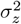. First, by the law of large numbers, and because the population is very large (*N* → ∞), the price in eq. (A-24) can be rewritten as

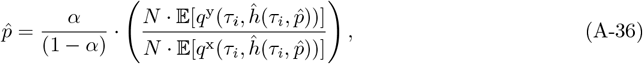

where the numerator and denominator represent expectations over the quantities produced and where we have used 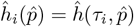 to make the trait dependence of individual *i* explicit. Second, if a random variable *X* has a small variance, then we can approximate the expectation of a function by the function of the expectation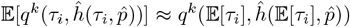. Furthermore, by definition, we have 𝔼 [*τ*_*i*_] = *θ*. Applying this approximation, eq. (A-36) can be rewritten as

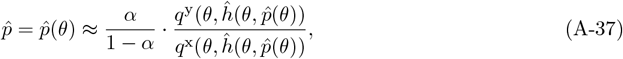

where 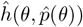 is the equilibrium time allocation of a resident individual. The last equation is equal to the equilibrium price in the mutant resident approximation when exchange takes place through a market, as in eq. (10). This shows that even if individuals exhibit small variations in their traits, the monomorphic approximation remains valid, as the price still converges to the same equilibrium value.

##### Appendix B.2.2 Explicit expression of equilibrium price and time allocation in a monomorphic resident population with market exchange

The equilibrium price in a mutant-resident population is explicitly defined by eqs. (9)–(10) when time allocation is fixed. However, when time allocation is a decision variable, these expressions become implicit because the price appears on both sides of the equation, requiring a numerical solution. While the equilibrium price under dyadic exchange can only be determined numerically, we show that for market exchange, a fully analytical expression for the price, and thus the equilibrium time allocations can be derived. This then allows for the evolutionary invasion analysis to be conducted without the need for numerical computations.

In a mutant-resident population, the price at equilibrium is given from eq. (10) by

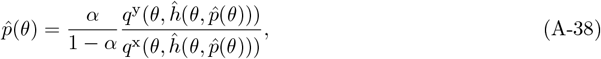

where 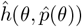 is the time allocation at equilibrium of a resident individual with trait *θ* facing an equilibrium price 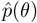. Substituting the functional forms of the quantities produced from eq. (1) into the last equation yields

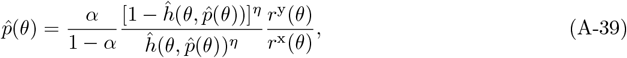

Substituting this last equation into the equation defining the equilibrium time allocation–where we adopt the formulation from eq. (A-15) for ease of derivation–and simplifying notation by writing 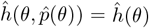, gives

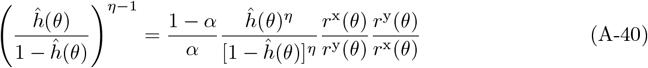

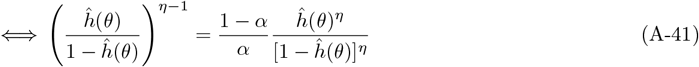

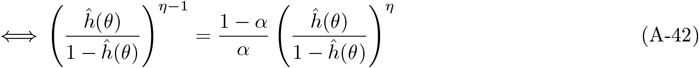

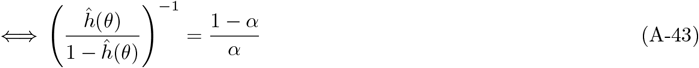

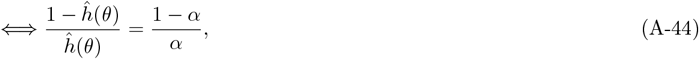

which produces

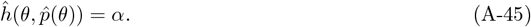

Hence, resident individuals allocate a proportion *α* of their time to produce good x. Substituting back the time allocation into eq. (A-39), the price simplifies to

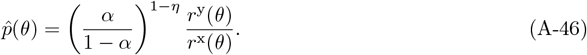

Substituting this price into eq. (11), the equilibrium time allocated to produce good × of a mutant is

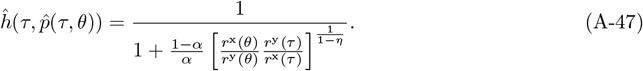

If we finally substitute the equilibrium price from eq. (A-46) and the time allocation from eq. (A-47) into the mutant payoff in eq. (6), it yields an explicit expression, which is used to conduct the invasion analysis

##### Appendix B.3 Evolutionary analysis with fixed time allocation

##### Appendix B.3.1 Autarky

We first carry out the evolutionary analysis under autarky (main text Section 3.1). Substituting the mutant payoff in eq. (12) into eq. (A-32), the payoff gradient for this case is

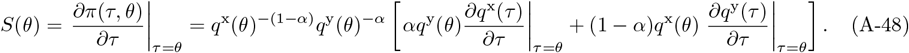

When the derivatives on the right-hand side have the same signs, the payoff gradient always point out in their direction and thus towards improving the production of both goods. When the derivatives on the right-hand side have opposite signs, there is a trade-off between the production of the two goods as a different value of the trait increases the production of one good but reduces the production of the other good. In this case, the terms in square brackets describe the sum of the gain or loss in the production of each good due to a change in the value of the trait, weighted by how much each good contributes to the payoff. As this weight depends of the quantities produced of the other good, this ultimately favours generalists who can produce both goods.

Setting *S*(*θ*) = 0 and rearranging shows that a singular point *θ*^*^ must satisfy

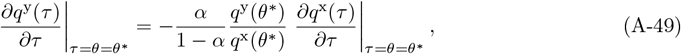

or equivalently eq. (13) of the main text.

Recall that that under autarky, the payoff function is assumed to admit a unique interior maximum with respect to the trait *τ*. Since the mutant payoff depends only on the mutant trait *π*(*τ, θ*) = *π*(*τ*), this assumption entails that the singular point corresponds to the trait value that maximises *π*(*τ*) and that the disruptive selection coefficient described by the second derivative 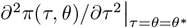 is negative at this point. Furthermore, it is straightforward to see that for the case where payoff depends only *τ*, the convergence stability coefficient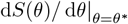, where *S*(*θ*) = *∂π*(*τ*)*/∂τ* |_*τ*=*θ*_, is equal to the disruptive selection coefficient:

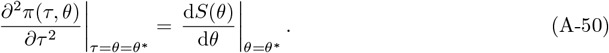

Therefore, the singular point is both convergent stable and uninvadable.

Substituting the production functions in eqs. (1)–(2) into eq. (A-49) and solving for *θ*^*^ yields eq. (14). Furthermore, at this point

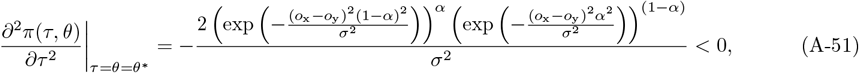

which confirms that the singular point is convergence stable and uninvadable under our specific production function.

##### Appendix B.3.2 Dyadic exchange

We now turn to the evolutionary analysis for the case of dyadic exchange (results of section 3.2). The mutant payoff in this section is the one defined in eq. (6) with price in eq. (9). Substituting it into eq. (A-32) but keeping the price in its symbolic form, the payoff gradient is

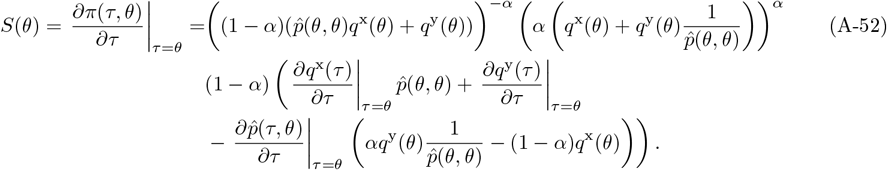

The last part of the expression can be rewritten using the definition of the quantity consumed of good × in eq. (A-21)

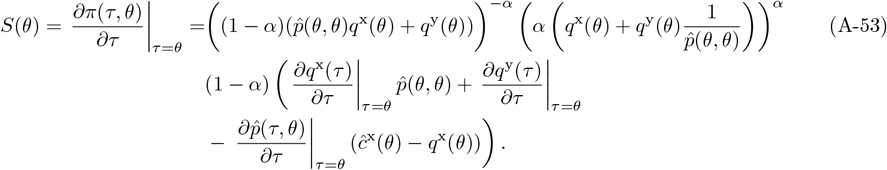

The part of the expression in the first line does not influence the sign of the payoff gradient. The rest of the expression can be decomposed in two parts. A change in the trait has again a direct cost and benefit by changing the quantities of goods produced. Here, the importance of a loss or gain in production of a good is captured by its price. Second, a change in the trait also has an indirect cost or benefit, described on the third line, as changing the quantities produced also modifies the price of goods. An increase in the price of good × is costly if the mutant is a net buyer of the good x, i.e. (*ĉ*^x^(*θ*) − *q*^x^(*θ*)) *>* 0 but beneficial if it is a net seller (*ĉ*^x^(*θ*) − *q*^x^(*θ*)) *<* 0.

Substituting the equilibrium price from eq. (9), evaluated in a resident population into eq. (A-53), the term in the third line vanishes

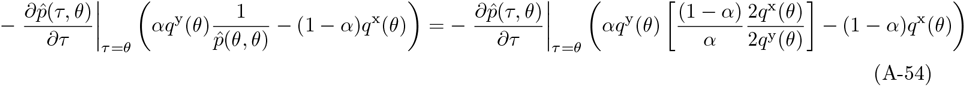

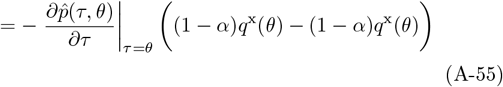

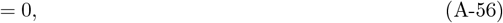

and the remaining expression is

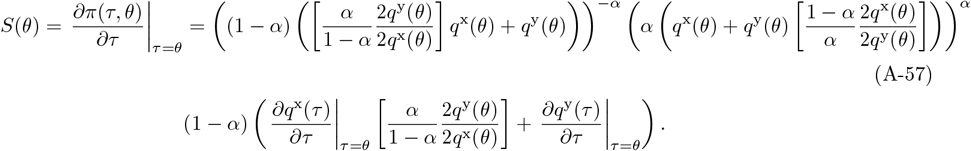

This simplifies as

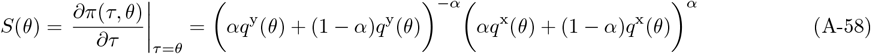

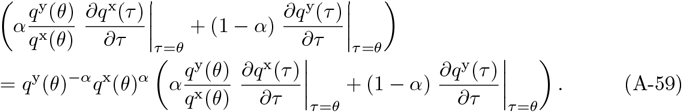

Factoring 1*/q*^x^(*θ*) from inside the parentheses out gives

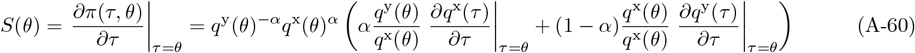

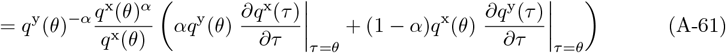

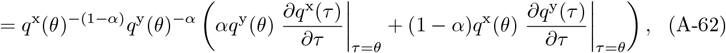

where the last line is equivalent to eq. (A-48), the payoff gradient without exchange.

Because the payoff gradient with dyadic exchange is equal to the one in autarky, it follows that the singular point and the convergence stability coefficient, which are both derived from the payoff gradient (recall Appendix B.1), remain the same than in autarky. By contrast, the interaction coefficient, which is null under autarky, is under the present exchange scenario given by

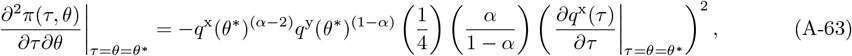

which yields eq. (15) of the main text and is always negative. Because of this and the observation that the right-hand of eq. (A-33) is conserved between dyadic exchange and autarky, a negative interaction coefficient implies an increase for the scope of disruptive selection, with the disruptive selection coefficient being given by

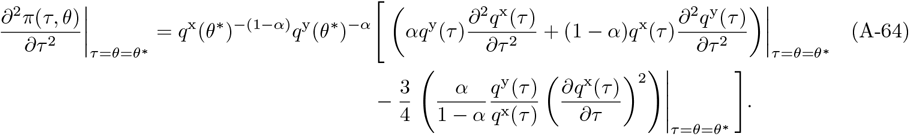

This expression can be rewritten by substituting one of the partial derivatives in the expression 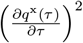 by using eq. (A-49), which leads to

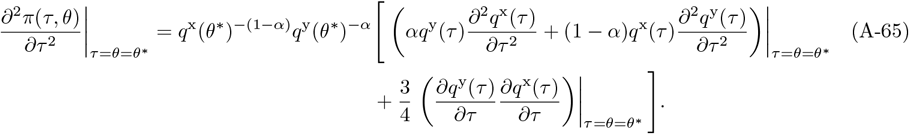

Identifying when this expression is positive and rearranging yields eq. (16) of the main text.

##### Appendix B.3.3 Market exchange

We here describe to the evolutionary analysis for the case of market exchange (results of section 3.3). In this case, the mutant payoff is the one described in eq. (6) with the price at equilibrium in eq. (10). Substituting this into eq. (A-32), the payoff gradient is given by

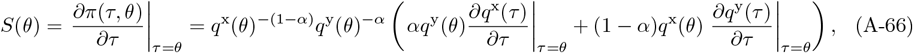

which is again equal to the one without exchange in eq. (A-48). Because the payoff gradient with market exchange is equal to the one in autarky, it follows that the singular point, which is the root of the payoff gradient, and the convergence coefficient 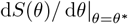 remain the same. Thus, there is a single convergent stable singular point.

Using again the mutant payoff under the present exchange scenario, we obtain the interaction coefficient

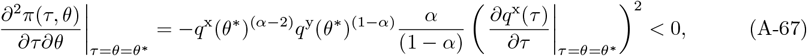

which is always negative and proportional to eq. (A-63), while the disruptive selection coefficient is

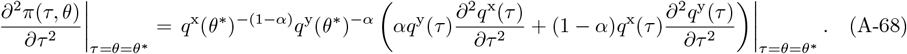

Identifying when this expression is positive yields

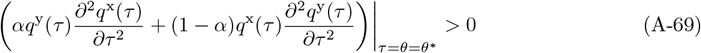

With market exchange, polymorphism emerges as long as the increase in production due to a deviation in the trait are greater than linear. This condition ensures that the gain in production in one good for a mutant is higher than its loss in the other good. Substituting the production function (eqs. (1)–(2)) yields eq. (18) of the main text.

#### Appendix B.4 Evolutionary analysis with time allocation decision

We now turn to the evolutionary invasion analysis when time allocation is a decision variable (results of section 4). While in the previous section, many results were expressed for arbitrary production functions, the situation is more complicated under the present scenarios, so the analytical results here below are presented only using the explicit production function eqs. (1)–(2).

##### Appendix B.4.1 Autarky

In autarky, the mutant payoff in eq. (6) simplifies to *π*(*τ, θ*) = [*q*^x^(*τ,ĥ* (*τ, θ*))]^*α*^[*q*^y^(*τ,ĥ* (*τ, θ*))]^1−*α*^, where we showed in Appendix A.2 that *ĥ* (*τ, θ*) = *α* and the payoff becomes a function of *τ* only. The payoff gradient derived from this expression yields

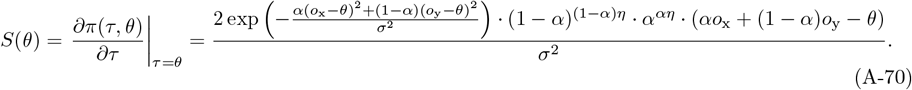

Setting to zero, and solving at *θ* = *θ*^*^, yields the singular point

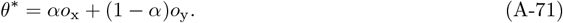

This is 0 for the parameters specified in the main text *α* = 0.5, *o*_x_ = −2 and *o*_y_ = 2.

Using the payoff gradient, we obtain

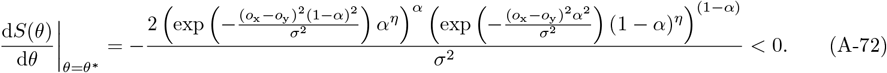

This expression is always negative, and thus the singular point is convergent stable and uninvadable (in force of eq. A-50).

##### Appendix B.4.2 Dyadic exchange

For the case of dyadic exchange, the price has no analytical expression and the evolutionary analysis is carried out numerically (see accompanying Mathematica notebook).

##### Appendix B.4.3 Market exchange

We now turn to the evolutionary analysis for the case of market exchange. Substituting the mutant payoff in eq. (6) with the price at equilibrium in eq. (A-46) and the time allocation in eq. (A-47) into eq. (A-32) gives the payoff gradient.

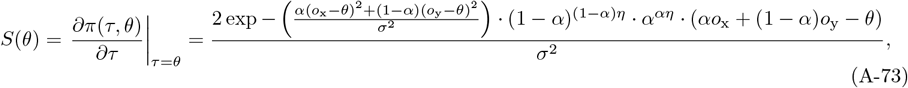

which is equal to the one without exchange given by eq. (A-70). Because the payoff gradient under market exchange is equal to the one in autarky, it follows that the singular point remains the same as given in eq. (A-71) and is convergence stable.

We obtain that the interaction coefficient is

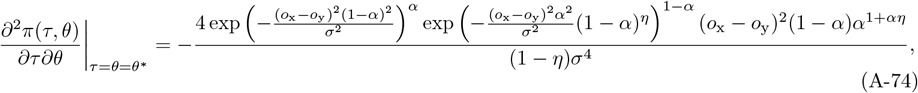

which is always negative, while

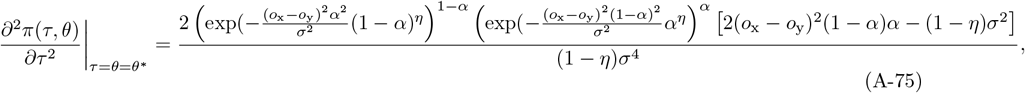

which is positive if the expression in the square brackets is positive. Rearranging this expression yields the condition given in the main text in eq. (19). Substituting the parameters used in the main text *α* = 0.5, *o*_x_ = −2 and *o*_y_ = 2, we obtain the condition for polymorphism to emerge used to generate the left panel of Fig. (2) as

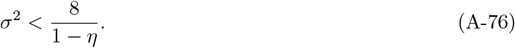

### Appendix C Single trait model: Simulations with asexual and sexual reproduction

We here describe the individual-based stochastic simulations we used to confirm our analytical results, explore more long-term evolution and further confirm that our findings also apply under sexual reproduction.

#### Appendix C.1 Simulation procedure

The simulations describe the evolution of a population of fixed and even size *N*. Each individual *i* ∈ *{*1, …, *N}* is characterised by its trait *τ*_*i*_ encoded at a single locus, and determined either by a single allele of value *a*_*i*_ under asexual reproduction or a pair of alleles of values (*a*_1*i*_, *a*_2*i*_) under sexual reproduction. The allelic values are defined on the same domain as the trait *a*_*i*_, *a*_1*i*_, *a*_2*i*_ ∈ ℝ. The initial population is a monomorphic population where all individuals have a trait of the same value.

The life cycle of individuals consists of discrete and non-overlapping generations as described in section 2.1, where the following events occur in cyclic order: (i) individuals are randomly paired with each other; (ii) time allocations are set (detailed below) (iii) individuals produce a quantity of each good as described in equation (1); (iv) individuals exchange quantities of goods and end up with quantities described in eqs. (A-21)–(A-20); (v) individuals reproduce following a Wright-Fisher process (see eq. (A-77) below) during which mutations occur; and (vi) adults of the previous generation perish. We repeat the procedure for a fixed number of generations.

##### Time allocation

In the version of the model where time allocation is fixed, the time allocation is automatically set to 0.5 for all individuals during step (ii). In the second version of the model where individuals choose their time allocation, the time allocation is given by eq. (A-19), where the price is found by solving eq. (A-24) numerically.

##### Mutation

For each allele, mutation occurs with probability *µ*_m_ and when a mutation occurs, a mutation effect drawn from a normal distribution centered on zero and with standard deviation *σ*_m_ is added to the allelic value.

##### Reproduction

During the reproduction stage, a new population of the same size *N* is created by sampling randomly with replacement in the parent population according to their payoff. The probability that an individual *i* with payoff *π*_*i*_ is the parent of an offspring is

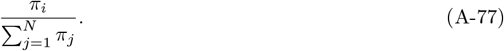

We carried out individual-based simulations under the above life-cycle for two genetic systems that differ in their mode of inheritance.

##### Asexual reproduction

Individuals carry a single allele determining the value of the trait *τ*_*i*_ = *a*_*i*_ for individual *i*. During reproduction, a single parent is sampled and the allelic value of the offspring is equal to the allelic value of this parent, subject to potential mutation.

##### Sexual reproduction

Individuals carry two alleles which, when averaged, determine the value of the trait *τ*_*i*_ = (*a*_1*i*_ + *a*_2*i*_)*/*2 for individual *i*. We assume here additivity of the allelic values following the continuum-of-alleles model framework [22, 23]. During reproduction, two parents are sampled (without replacement) and the offspring inherits one randomly sampled allele from each parents.

#### Appendix C.2 Results

##### Asexual reproduction

All simulation results for the asexual model are presented and discussed in the main text.

##### Sexual reproduction

Considering sexual reproduction, we first run simulations with fixed parameter values assuming that exchange is present from the start. We restrict the analysis to fixed time allocation in order to focus solely on how sexual reproduction affects the evolutionary outcomes. In the presence of exchange, the population initially evolves towards a generalist phenotype with a trait value of 0, between the two production optima (Figure S1). As in the asexual model, exchange subsequently promotes genetic diversity and the emergence of distinct morphs.

Sexual reproduction, however, leads to different long-term patterns than under asexual reproduction. While selection favours specialised individuals with trait values near one of the optima, mating between individuals with different trait values produces offspring with intermediate, generalist phenotypes, through allele segregation. Although these intermediate phenotypes are disfavoured by selection and eventually eliminated, they are continuously reintroduced each generation through recombination. As a result, selection under sexual reproduction maintains 3 different morphs in the population.

To further confirm that genetic diversity in the population is indeed markedly increased as a result of introducing exchange under sexual reproduction, we evaluated the variance 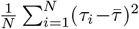 in the genetic trait in the population, where 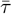 is the average trait value, over time across 10 simulations replicates and compared the results with the asexual reproduction case (Figure S2). As described in the main text, the variance consistently increases with time after the branching point is reached, as predicted by the asexual model (see Appendix B.1 for details). The genetic variance is smaller with sexual reproduction as offspring of specialised individuals reintroduce unfit generalist individuals. Overall, these results confirm that exchange promotes and maintains genetic diversity under sexual reproduction.

**Figure S1.**
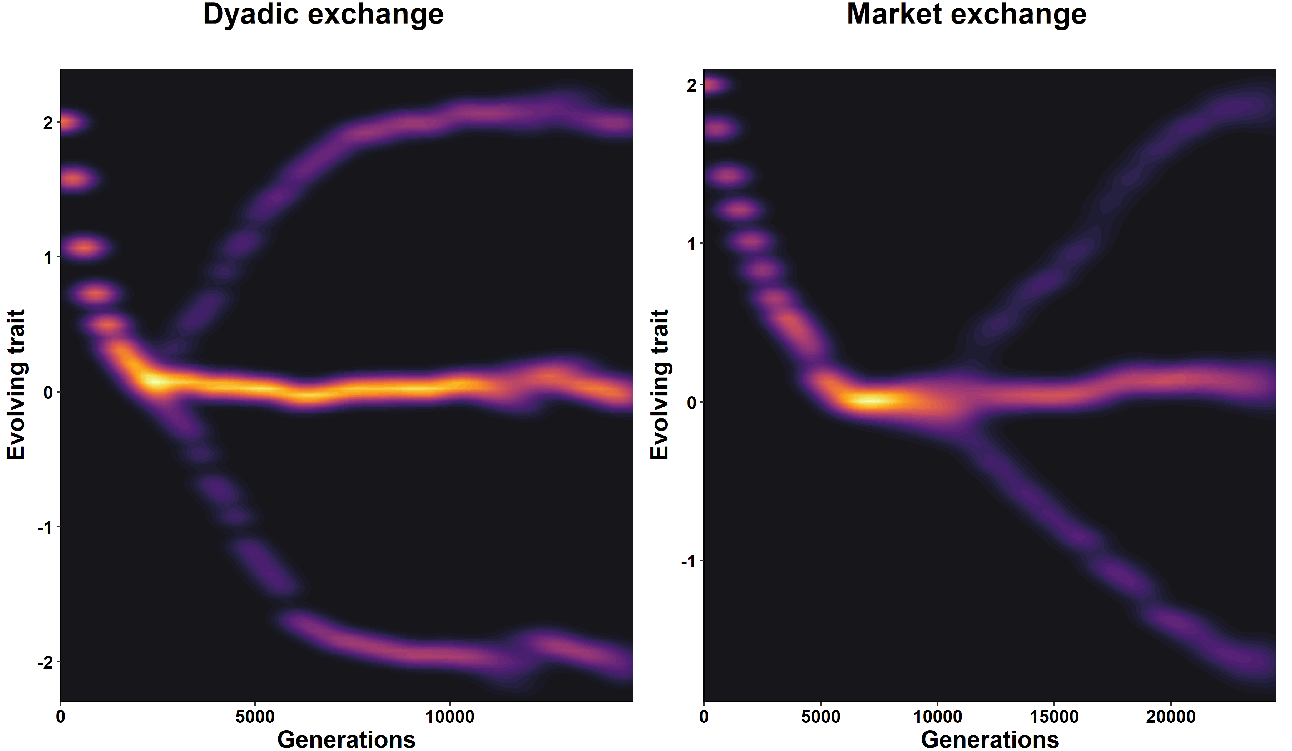
Distribution of trait values over generations under dyadic exchange and market exchange when the trait is encoded by a single diploid locus. Each allele effect is initially set to 2. Parameters: equal value of goods *α* = 0.5, optima of production *o*_x_ = −2, *o*_y_ = 2, population size *N* = 5000, production breadth *σ*^2^ = 1 for dyadic exchange and *σ*^2^ = 5 for market exchange, mutation parameters *µ*_m_ = 0.01, *σ*_m_ = 0.02.

**Figure S2.**
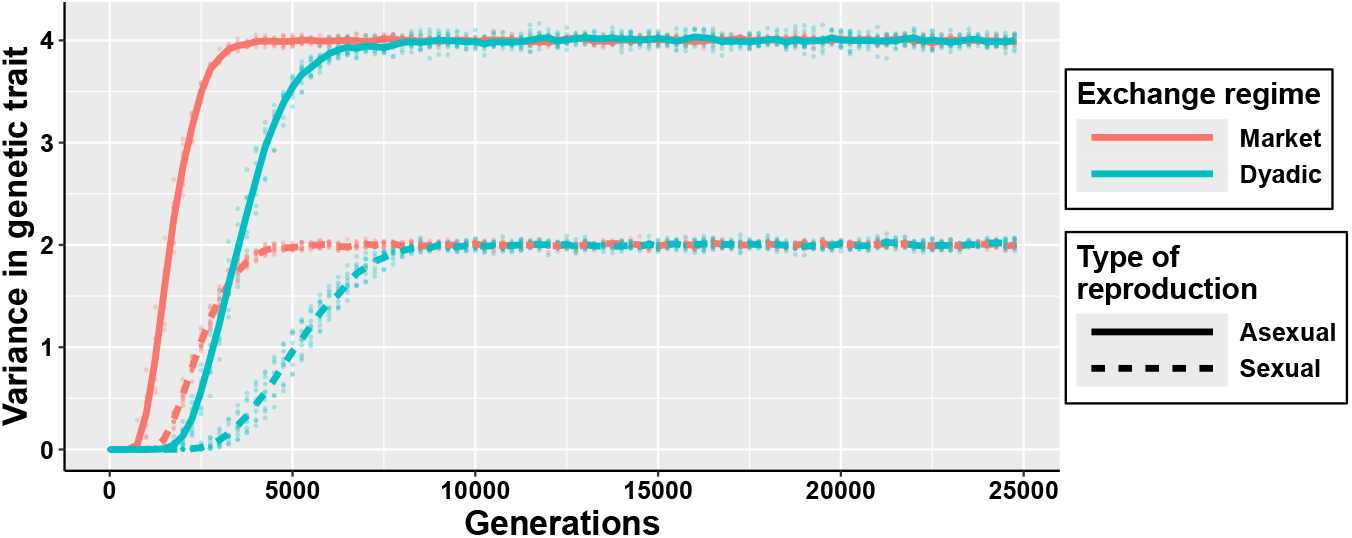
Variance in the genetic trait over generations under different exchange regimes (dyadic or market) and reproduction modes (asexual or sexual). Dots show values from 10 simulation replicates; lines indicate the mean variance. Variance is computed as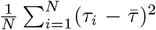 where 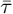 is the average population trait value. Initial trait value: 2. Parameters: production breadth *σ*^2^ = 1, equal value of goods *α* = 0.5, optima of production *o*_x_ = −2, *o*_y_ = 2, population size *N* = 5000, mutation parameters *µ*_m_ = 0.01, *σ*_m_ = 0.02.

### Appendix D Two traits model: Model specification and simulations

In the model of the main text, we considered a single trait that simultaneously affects the ability to produce both goods, which might suggest that exchange promotes genetic diversity only for this kind of trait. To assess the generality of our results, we examine an alternative model in which the abilities to produce each good depend on distinct traits, given that there is limited resources that can be allocated to develop both these traits. For clarity, the key differences between the alternative versions of the model are summarised in Table S1.

#### Appendix D.1 Model specification

The abilities to produce each good now depend on distinct traits, denoted *τ* ^x^ ∈ [0, 1] and *τ* ^y^ ∈ [0, 1]. In contrast to the main single-trait model, where production depends on the proximity of the trait value to an optimum, a higher value of a trait here always increases the quantity produced, such that an individual with traits 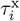 and 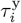 produce quantities

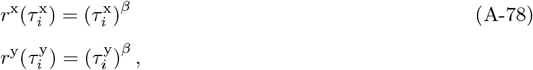

where *β >* 0 captures the trait elasticity of scale, quantifying returns to scale from investing in each trait. In this section, we restrict the analysis to fixed time allocation and thus the quantities produced depend only on the traits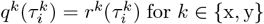. To reflect that a limited amount of resources is available for the development of these traits, we assume that the sum of the two trait values is bounded 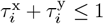 for individual *i*.

To better understand how the two-trait model relates to the single-trait model, we illustrate the space of possible production outcomes in Figure S3. These are the possible pairs of quantities of goods × and y that an individual can produce given the possible trait values it can bear. The solid lines in both panels represent the trait production-possibility frontiers (or trade-off curves), which give, among all possible trait combinations an individual can bear, the maximum production level *q*^y^ of good y that this individual can produce for any given production level *q*^x^ of good x. In the single trait model, the production-possibility frontiers is obtained as the set of production pairs (*q*^x^(*τ*), *q*^y^(*τ*)) by varying *τ* through some interval (that is [−10, 10] in Figure S3). Under the single-trait model, evolution is constraint to take place on this production-possibility frontier. In the two trait model, the production-possibility frontier is obtained as the set of production pairs (*q*^x^(*τ* ^x^), *q*^y^(*τ* ^y^)) obtained by making the constraint 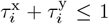binding, whereby *τ* ^x^ = 1 − *τ* ^y^, and then varying *τ* ^y^ through the interval [0, 1]. In contrast to the single trait case, in the two-trait model individuals are allowed to evolve within the entire feasible space below the trait production-possibility frontier (represented by the shaded area for the case *β* = 2 in Figure S3), where the total investment may fall short of the maximum resource budget. We note that it is customary in evolutionary biology to restrict trait evolution to the trait production-possibility frontier, as this simplifies the analysis without loss of qualitative generality. Below the trait production-possibility the constraints are not binding (so an individual is wasting resources), meaning that evolution necessarily selects for the trait values to be on the trait production-possibility frontier.

**Figure S3.**
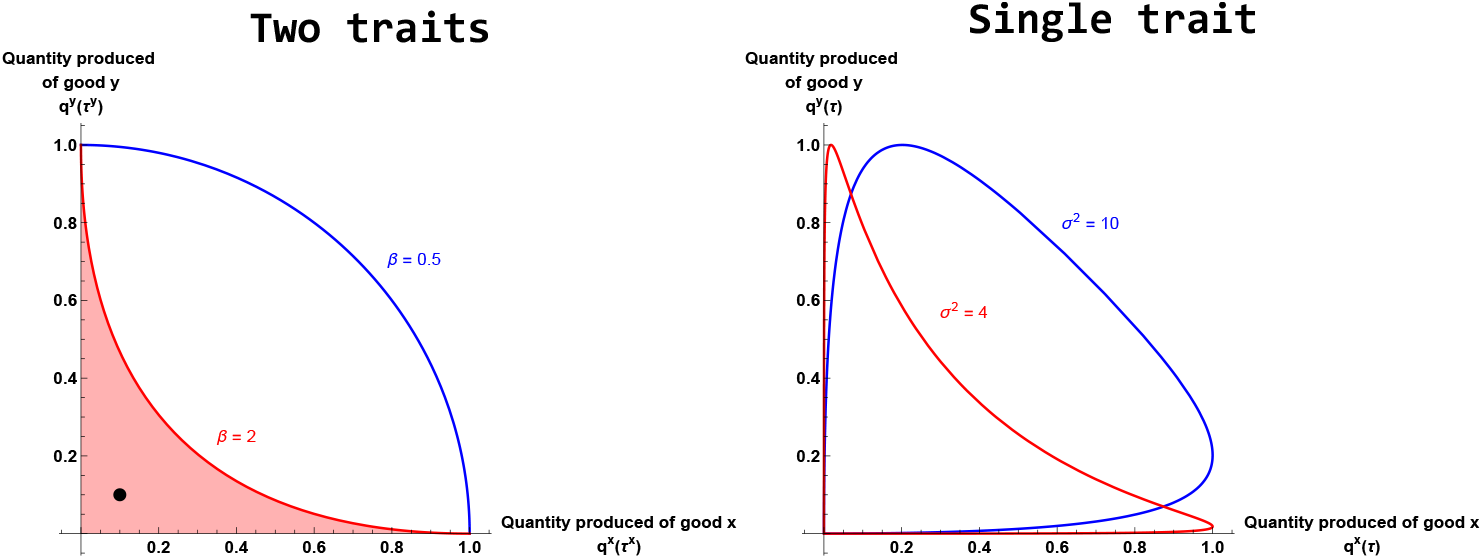
Quantities of good × and y produced in the single-trait and two-trait models, for a case where polymorphism can occur (*σ*^2^ = 4, *o*_x_ = −2, *o*_y_ = 2 for single trait, and *β* = 2 for two traits) and where it cannot (*σ*^2^ = 10, *o*_x_ = −2, *o*_y_ = 2, for single trait and *β* = 0.5 for two two trait). The solid lines in both panels represent the trait production-possibility frontiers (or trade-off curve) and polymorphism can occur only if the production-possibility frontier is a convex curve for some region in the (evolving) trait space. In the convex region, and given some initial production pair (*q*^x^, *q*^y^), an individual producing one additional unit of good × results in a decrease in its production of good y that is falling as more × is initially produced. This tends to favour specialisation. By contrast, if the production-possibility frontier is a concave curve, then given some initial production pair (*q*^x^, *q*^y^), an individual producing one additional unit of good × results in a decrease of its production of good y that is increasing as more × is initially produced. This tends to inhibit specialisation. In the single-trait model, individuals always lie on the trait production-possibility frontier, using their entire resource budget. In the two-trait model, however, individuals may evolve within the full feasible region below the frontier (shaded in red for *β* = 2), where trait values may sum to less than one. In the simulations, individuals in the initial population have traits values of 0.1 for each trait, represented by the black dot.

We conduct simulations considering trait elasticity of scale *β* = 5 for dyadic exchange and *β* = 2 for market exchange. This choice reflects the conditions derived in eqs. (16) and (A-69), where the emergence of genetic diversity requires sufficiently large second derivatives of the production functions. Under the present production function in eq. (A-78), this condition is satisfied when *β* ≫ 1, corresponding to the production-possibility frontier being a convex curve (see Figure S3).

#### Appendix D.2 Simulation procedure

The general setting of the simulations for the two-trait model are similar to that of the one-trait model described in Appendix C.1, except that individual *i* ∈ *{*1, …, *N}* is now characterised by two traits 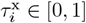 and 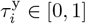 encoded at two distinct loci, by either a single allele of value 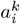 under asexual reproduction or a pair of alleles of values 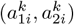 for *k* ∈ *{*x, y*}* under sexual reproduction. The allelic values are defined on the same domain than the trait [0, 1]. The initial population is a monomorphic population, with all individuals having allelic values set to 0.1, yielding *τ* ^x^ = *τ* ^y^ = 0.1.

The two genetic systems we consider differ in their mode of inheritance as follows.

##### Asexual reproduction

During reproduction, a single parent is sampled and the allelic values of the offspring are equal to the allelic values of this parent, subject to potential mutation. During development, the trait values of the offspring are set equal to the allelic values 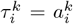 for *k* ∈ *{*x, y*}* of the corresponding locus if they satisfy the constraint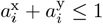, and rescaled proportionally otherwise:

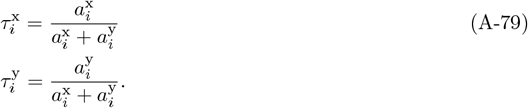

This procedure preserves the relative investment between traits while ensuring that individuals remain within the constraint. It reflects a biological scenario where genes specify target trait values, but realised trait values are shaped jointly by genetic targets and resource limitations.

##### Sexual reproduction

During reproduction, two parents are sampled (without replacement). The offspring inherits one randomly sampled allele from each parents at each locus, assuming free recombination. During development, trait values are set to the average of the allelic values at the corresponding locus 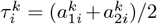 for *k* ∈ {x, y} if they satisfy the constraint 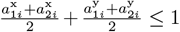, and are rescaled proportionally otherwise

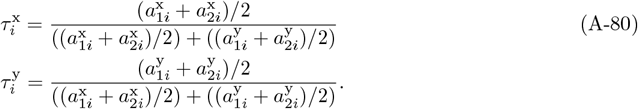

#### Appendix D.3 Results

We now present the simulation results for the two-trait model.

##### Appendix D.3.1 Asexual reproduction

Under both dyadic and market exchange (Figure S4), populations starting with low trait values first converge toward trait values around 0.5. At this point, individuals fully invest their available resources and divide them evenly between the two traits. Subsequently, the two traits diverge, with individuals evolving towards specialisation, expressing either high values of *τ* ^x^ and low values of *τ* ^y^, or *vice versa*. On the long term, selection maintains two specialised morphs. Thus, exchange promotes genetic diversity even when traits are encoded by distinct genes, provided individuals face a resource constraint and trait investment yields sufficiently strong returns.

##### Appendix D.3.2 Sexual reproduction

We now examine if our findings of the previous section are robust under sexual reproduction. Simulations show the same initial pattern, with a convergence of both traits towards 0.5 and then a strong increase in genetic diversity (S5).

**Figure S4.**
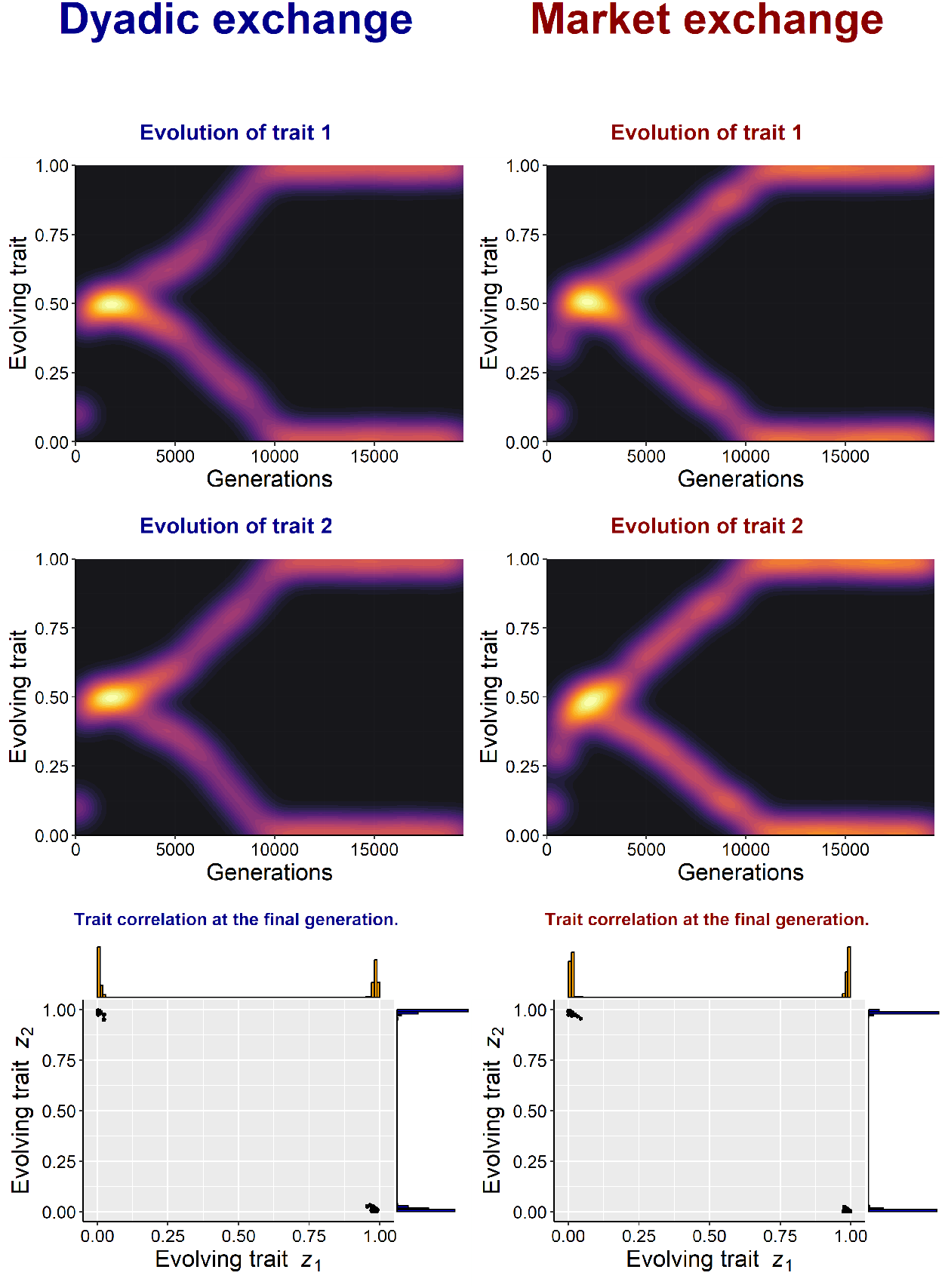
Distribution of the two trait values as a function of generations under dyadic exchange (left) and market exchange (right), assuming asexual reproduction. The bottom panels show the joint distribution of two evolving traits at the final generation. Points represent individuals and densities show the distribution of each trait across the population. The initial values of each trait are 0.1. Parameters: trait elasticity of scale *β* = 5 for dyadic exchange and *β* = 2 for market exchange, equal value of goods *α* = 0.5, population size *N* = 5000, mutation parameters *µ*_m_ = 0.01, *σ*_m_ = 0.005.

**Figure S5.**
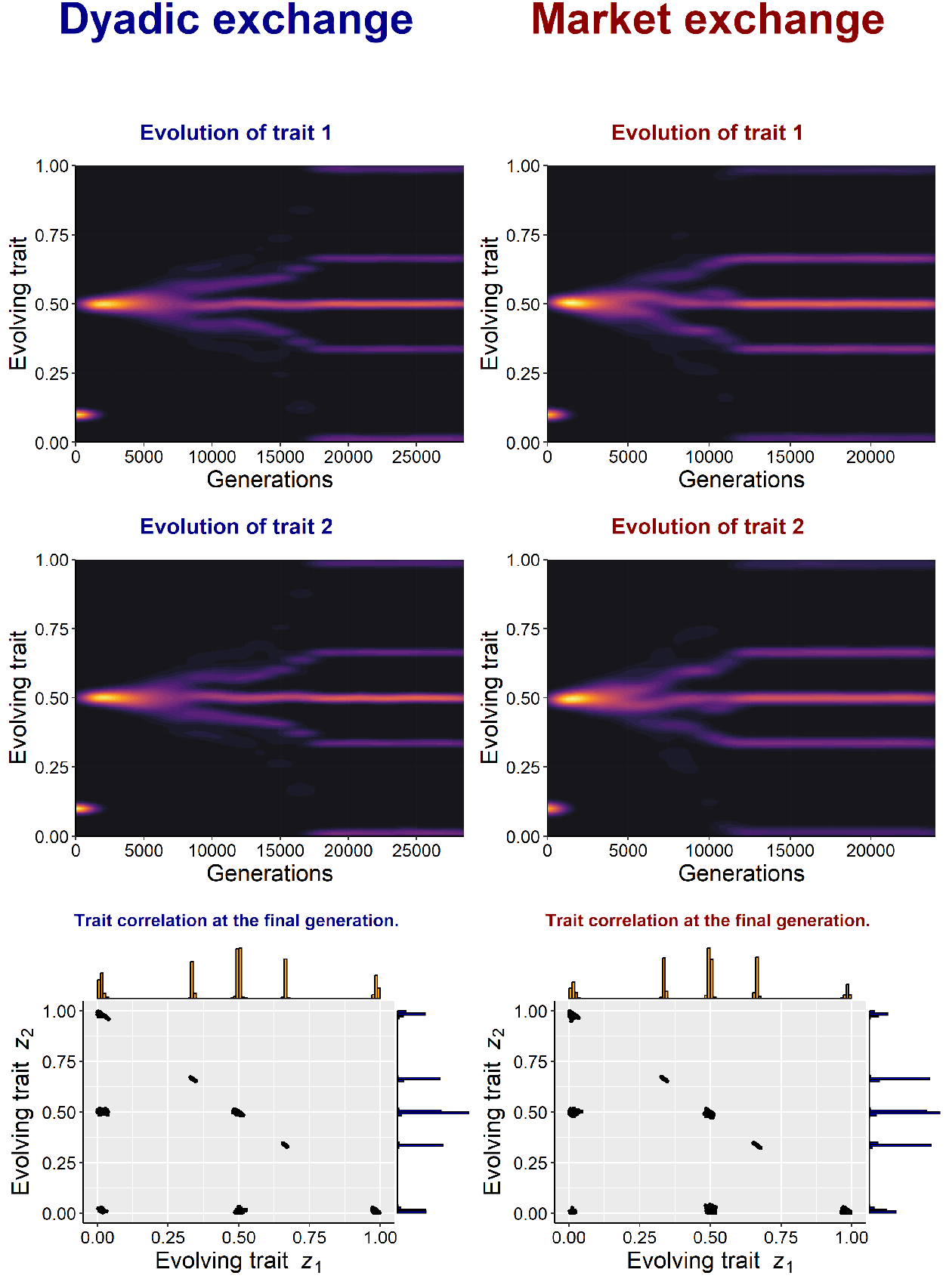
Distribution of the two trait values over generations under dyadic exchange (left) and market exchange (right), assuming sexual reproduction. The bottom panels show the joint distribution of two evolving traits at the final generation. Points represent individuals and densities show the distribution of each trait across the population. The initial values of each trait are 0.1. Parameters: trait elasticity of scale *β* = 5 for dyadic exchange and *β* = 2 for market exchange, equal value of goods *α* = 0.5, population size *N* = 5000, mutation parameters *µ*_m_ = 0.01, *σ*_m_ = 0.01.

On the long term, we observe the emergence of five distinct phenotype per trait, or eight distinct morphs when considering trait combinations across both loci. This pattern arises because of the interaction between selection favouring extreme morphs and recombination and segregation mixing them. Specifically, selection favours individuals with trait values close to either 0 or 1, which in turn selects for alleles with effects of either 0 or 1 at both loci. In other words, at equilibrium, each allele effectively behaves as a discrete unit taking value 0 or 1. Recombination and segregation generates a range of allele combinations. After applying possible renormalisation due to the resource constraint, the realised trait values include 0, 0.33, 0.5, 0.66, and 1 for each trait. For instance, an individual carrying allele pairs (1, 1) and (1, 0) for the two loci would have as phenotype *τ* ^x^ = 2*/*3 and *τ* ^y^ = 1*/*3. Overall, this mechanism produces a maximum of 3 *×* 3 = 9 phenotypes before normalisation. However, due to redundancy after rescaling (the genotype composed entirely of alleles with value 1 produces the same phenotype *τ* ^x^ = 0.5 and *τ* ^y^ = 0.5 after normalisation than the genotype that is fully heterozygous at both loci), only eight distinct phenotypes are ultimately observed in the population (observed in bottom panel of Figure S5).

### Appendix E Summary of the different model specifications

**Table S1.**
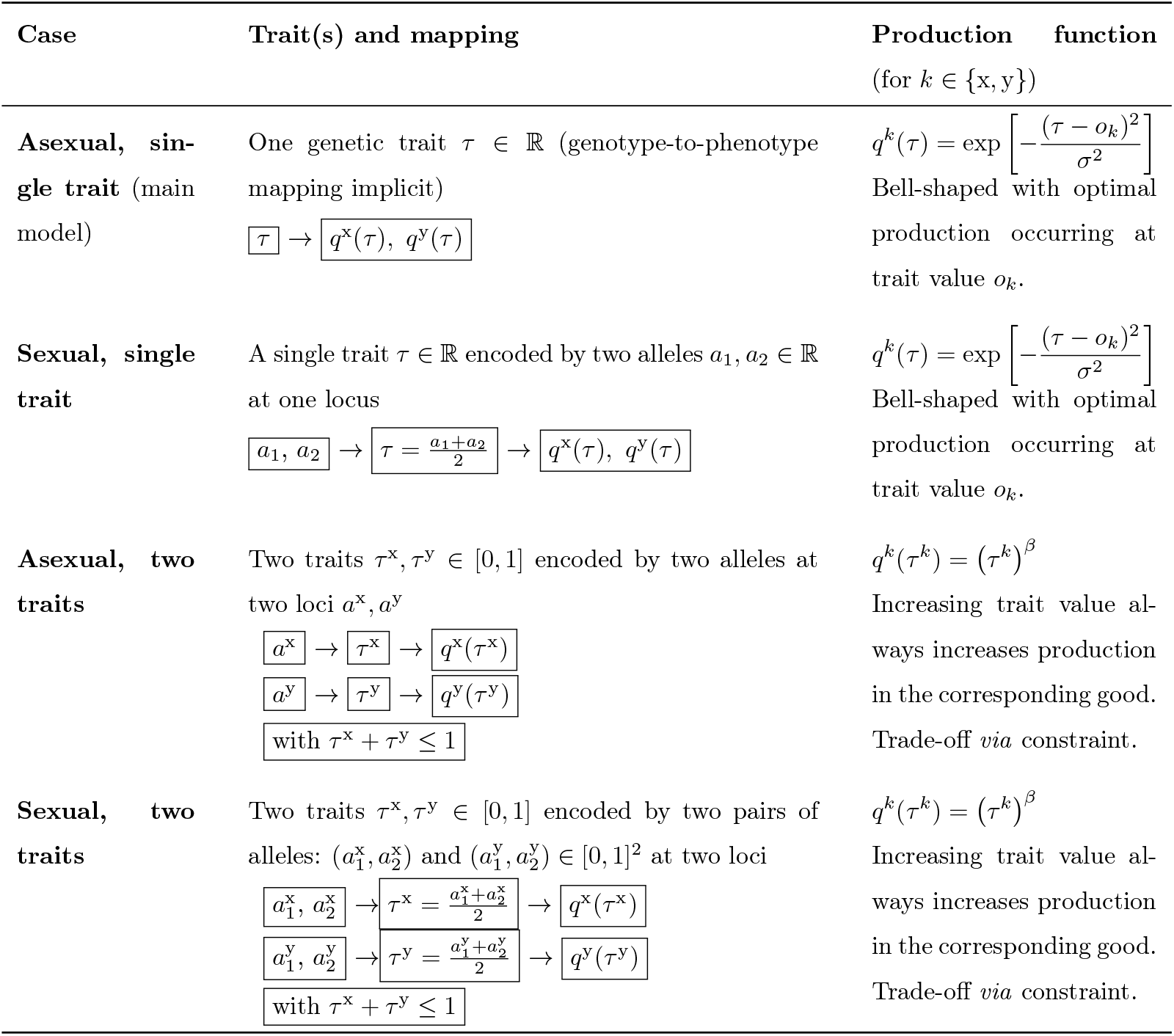
Summary of the traits, their encoding, and their mapping to production across the alternative versions of the model considered. In each case considered, genetic diversity emerges when exchange is present and following the predictions of the analytical model. This confirms that the main results are robust to changes in reproduction mode and genetic architecture.

## Supplementary Figures

**Figure S6.**
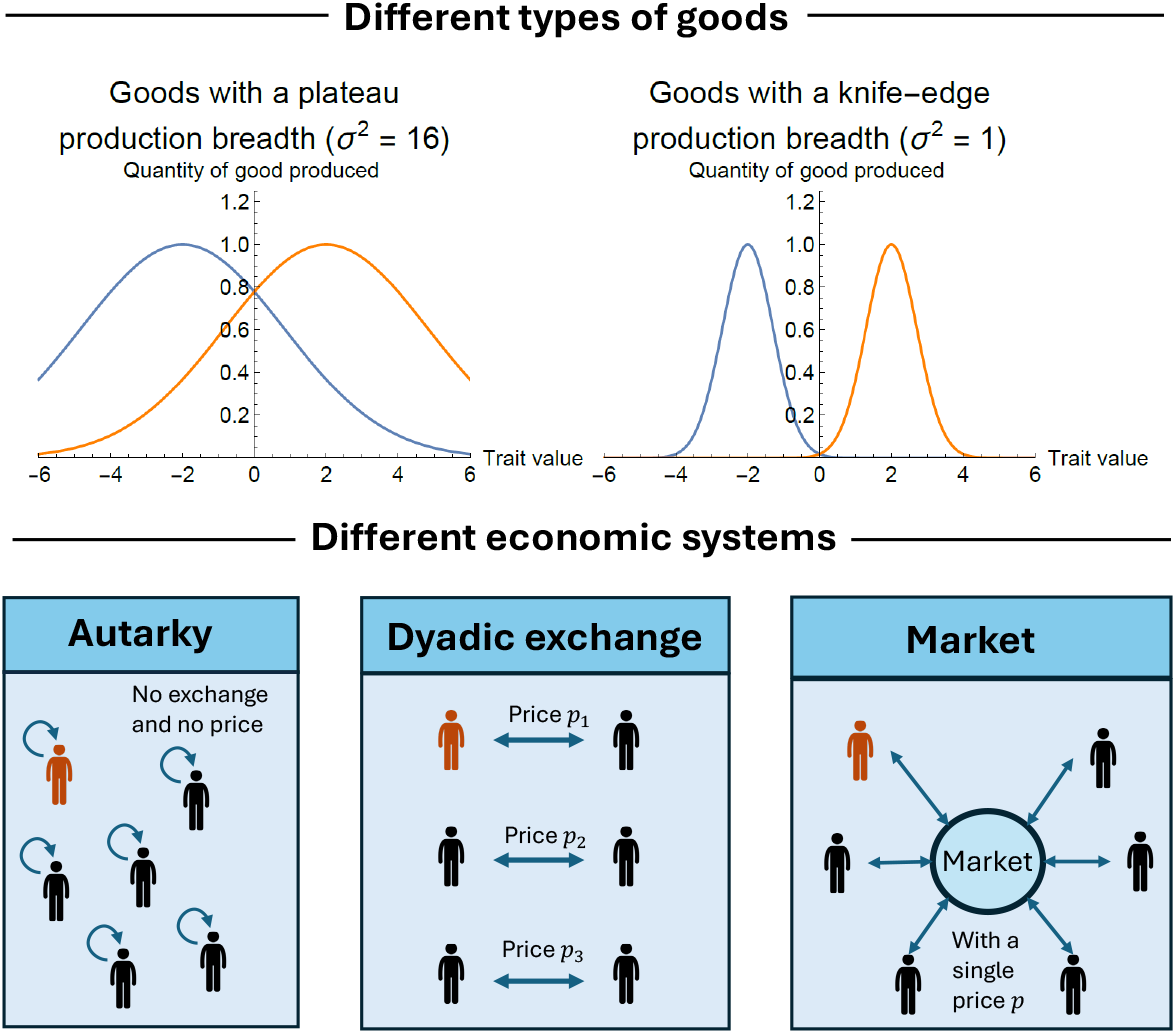
(Top) Quantity of good *x* and *y* produced, namely eq. (2), as a function of trait value, assuming an equal time allocated to the production of each good. The plots illustrate two types of goods, differing in production breadth *σ*^2^, with either high or low values. The optima of production of each good are *o*_x_ = −2 and *o*_y_ = 2. (Bottom) Modes of exchange considered: (i) autarky where exchange is not possible, (ii) dyadic exchange, where individuals exchange in randomly formed pairs, with each pair potentially settling on different prices independently, and (iii) market, where a single price is determined by aggregate production across all individuals.

**Figure S7.**
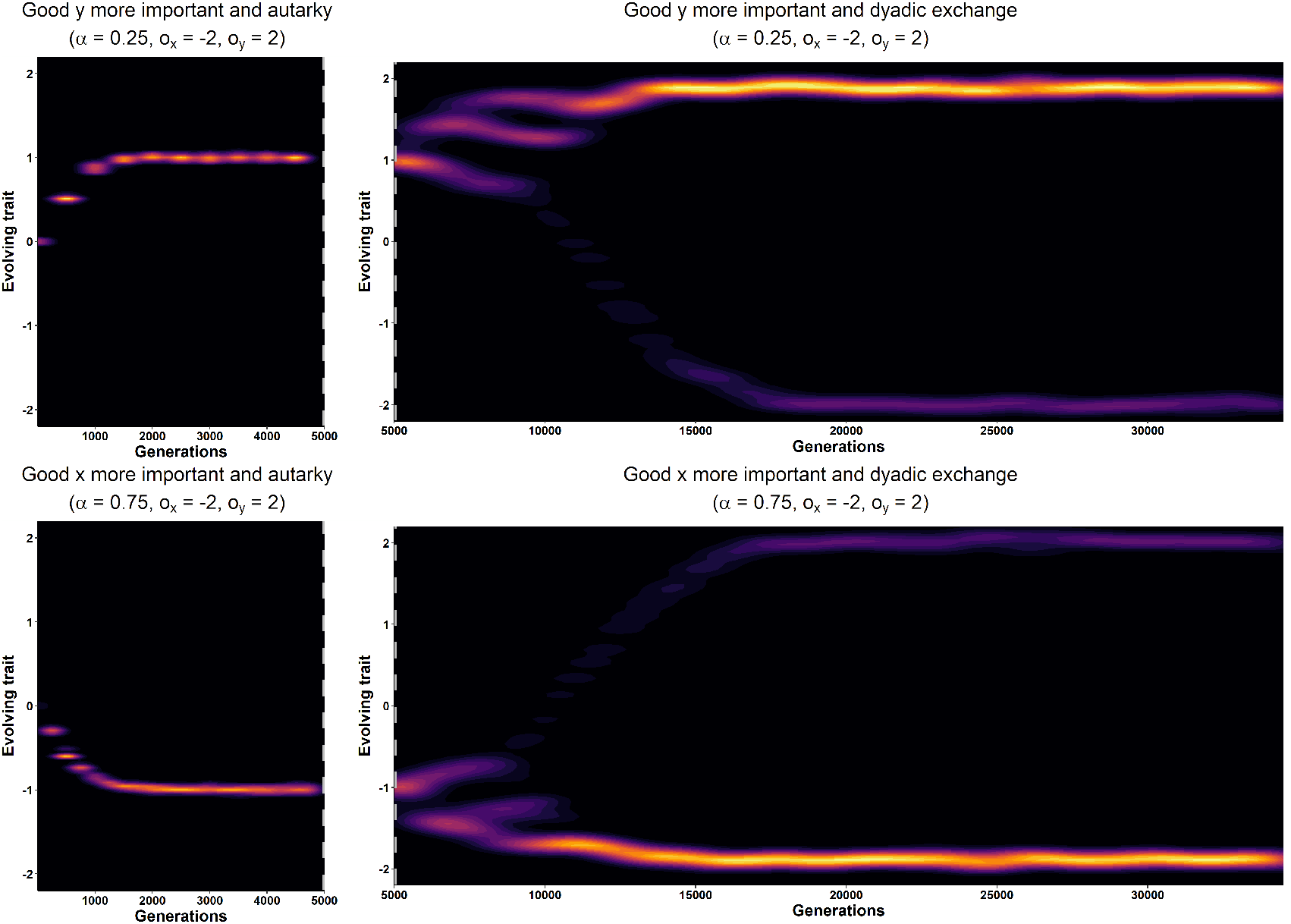
Evolution of the distribution of trait over generations when goods have different values and for fixed time allocation. The top panel represents a scenario where good y is more important than good × (*α* = 0.25), while the bottom panel represents the opposite case, where good × is more important than good y (*α* = 0.75). The population remains in autarky for the first 5000 generations, after which dyadic exchange is introduced (indicated by the grey dotted line). For better visualization of the density distributions, the plots are split into two phases: the autarky period and the period with dyadic exchange. Parameters: production breadth *σ*^2^ = 1, optima of production of each good *o*_x_ = −2 and *o*_y_ = 2, population size *N* = 5000, mutation parameters *µ*_m_ = 0.01, *σ*_m_ = 0.02.

**Figure S8.**
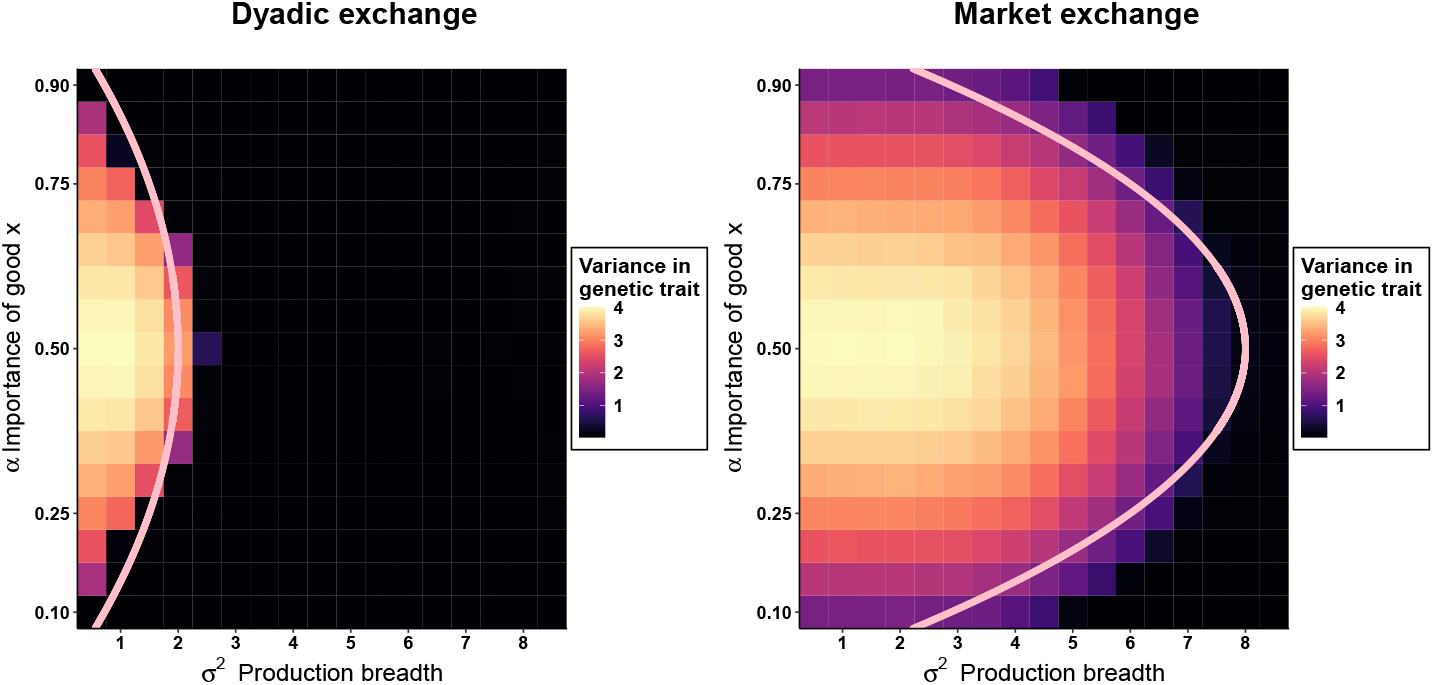
Variance in genetic trait as a function of the production breadth *σ*^2^ and relative value of good × (*α*) for two exchange regimes, dyadic exchange and market exchange, when effects of mutation are large *σ*_m_ = 0.2. The SD of mutations is 10 times higher than in the baseline scenario. The variance is calculated as 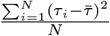 where 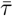 is the average trait value. Values are measured after 10000 generations and averaged across 10 replicates. The pink lines represent the combination of parameters for which genetic diversity is predicted to to emerge based on eqs. (17) and (18). Parameters: optima of production *o*_x_ = −2, *o*_y_ = 2, population size *N* = 5000, mutation rate *µ*_m_ = 0.01. Note that small variance in trait values can also arise outside the parameter range predicting the emergence of polymorphism, since large mutations generate diversity in the trait and that this diversity is always temporarily favoured, as shown in eqs. (15) and (A-67).

**Figure S9.**
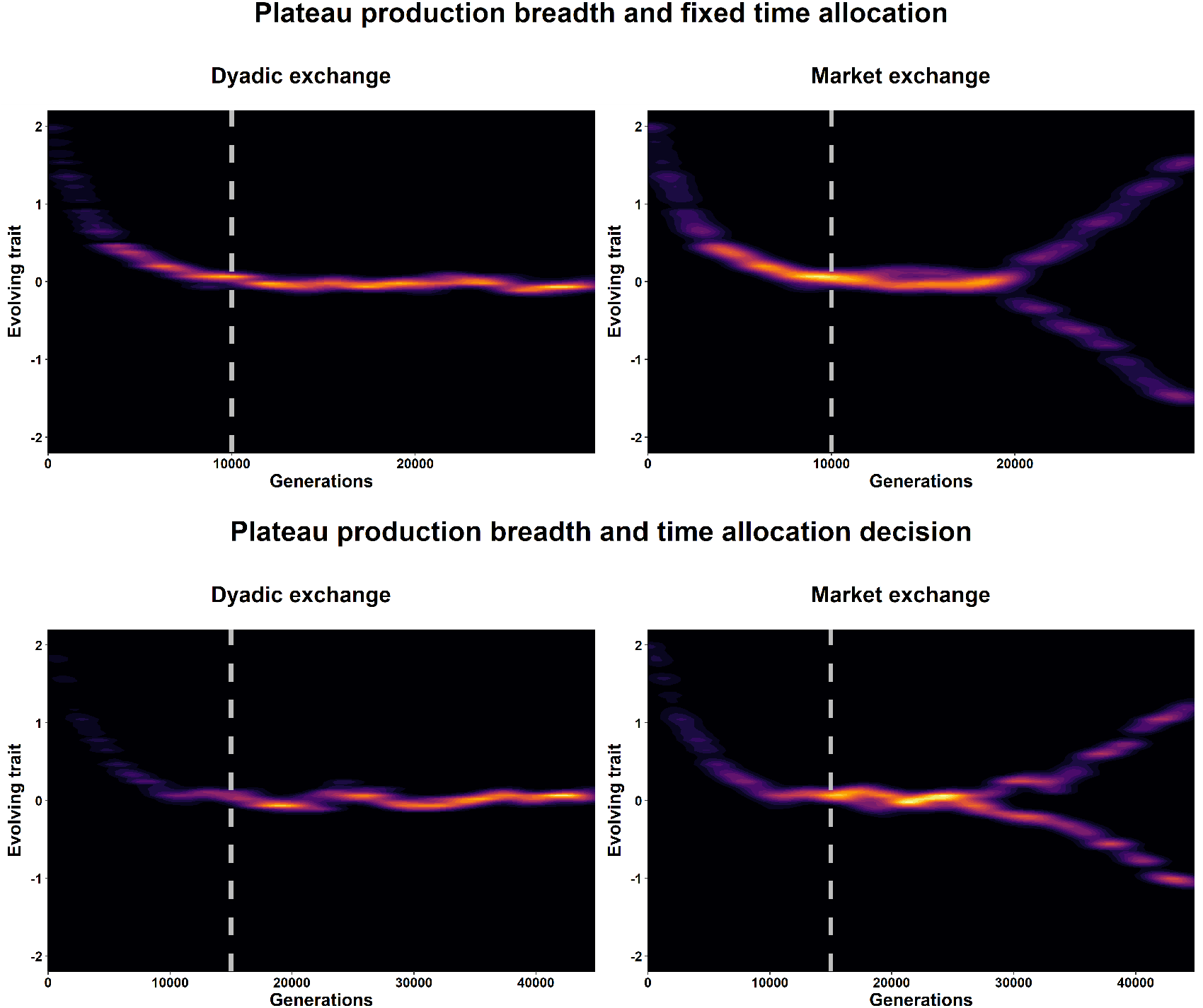
Evolution of the distribution of the trait over generations under two exchange regimes: dyadic exchange, where exchange occurs between isolated pairs of individuals, and market exchange, where all individuals interact through a common market. The goal is to illustrate how market exchange creates conditions more conducive to the emergence of polymorphism than dyadic exchange. To highlight this effect, we select high values of production breadth for which our theoretical results predict the emergence of polymorphism under market exchange but not under dyadic exchange: specifically, *σ*^2^ = 5 for fixed time allocation and *σ*^2^ = 10 when time allocation is a decision. Exchange is initially absent and is introduced only after 10000 or 15000 generations (indicated by the grey dotted line). Parameters: optima of production of each good *o*_x_ = −2 and *o*_y_ = 2, equal value of goods *α* = 0.5, population size *N* = 1000, mutation parameters *µ*_m_ = 0.01, *σ*_m_ = 0.02.

**Figure S10.**
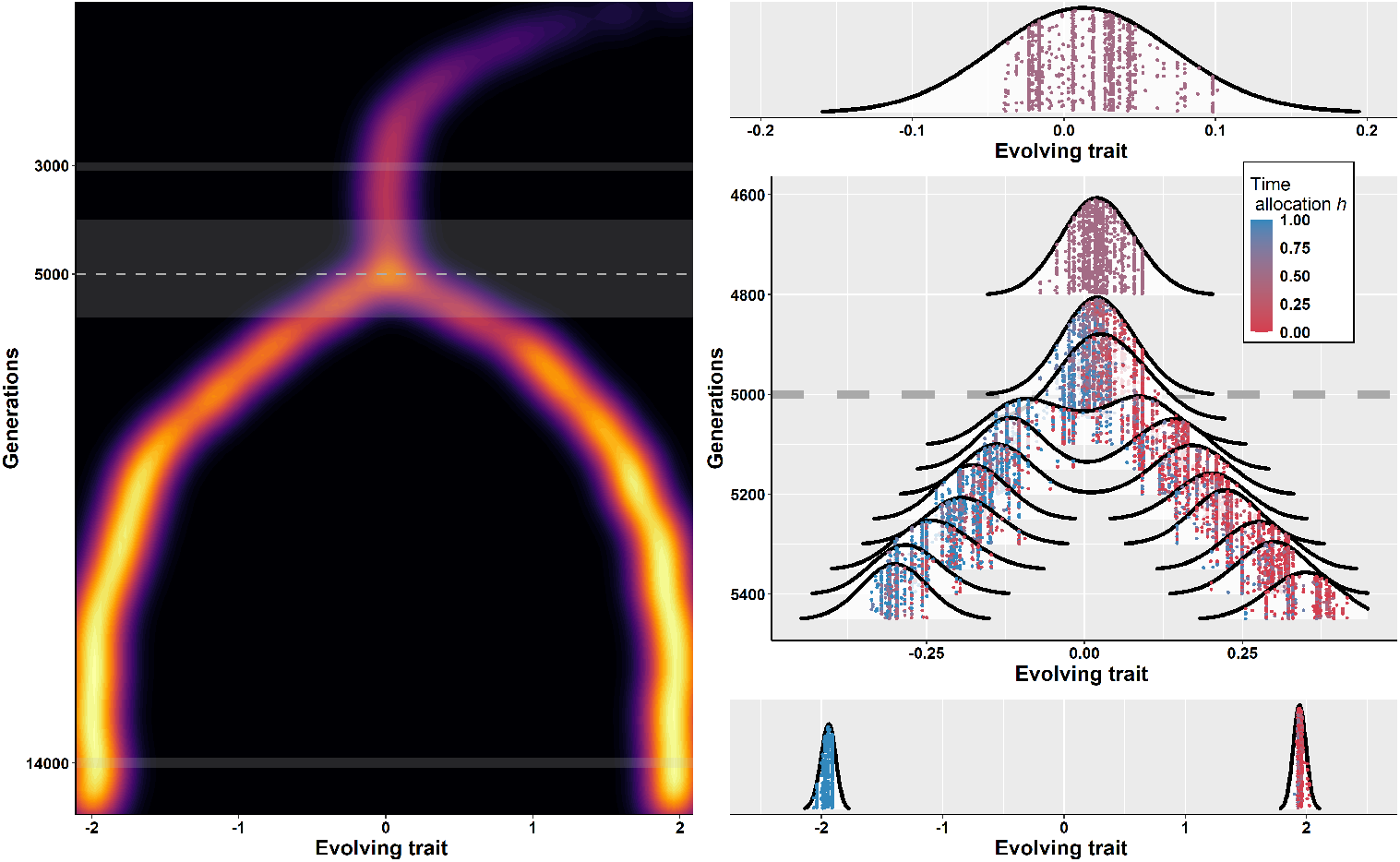
Evolution of genetic diversity and economic specialisation. (Left) Evolution of trait over generations. The population remains in autarky for the first 5000 generations, after which dyadic exchange is introduced (indicated by the grey dotted line). (Right) Distribution of trait (x-axis) and time allocation (colour). Each point represents a single individual. The plot is divided in three parts, each focusing on a particular time period, (top) at 3000 generations when exchange is absent, (middle) at 5000 generations when exchange is introduced and (bottom) at 14000 generations to show the long-run outcome. Parameters: elasticity of scale *η* = 0.9, production breadth *σ*^2^ = 1, equal value of goods *α* = 0.5, optima of production *o*_x_ = −2, *o*_y_ = 2, population size *N* = 1000, mutation parameters *µ*_m_ = 0.01, *σ*_m_ = 0.02.

**Figure S11.**
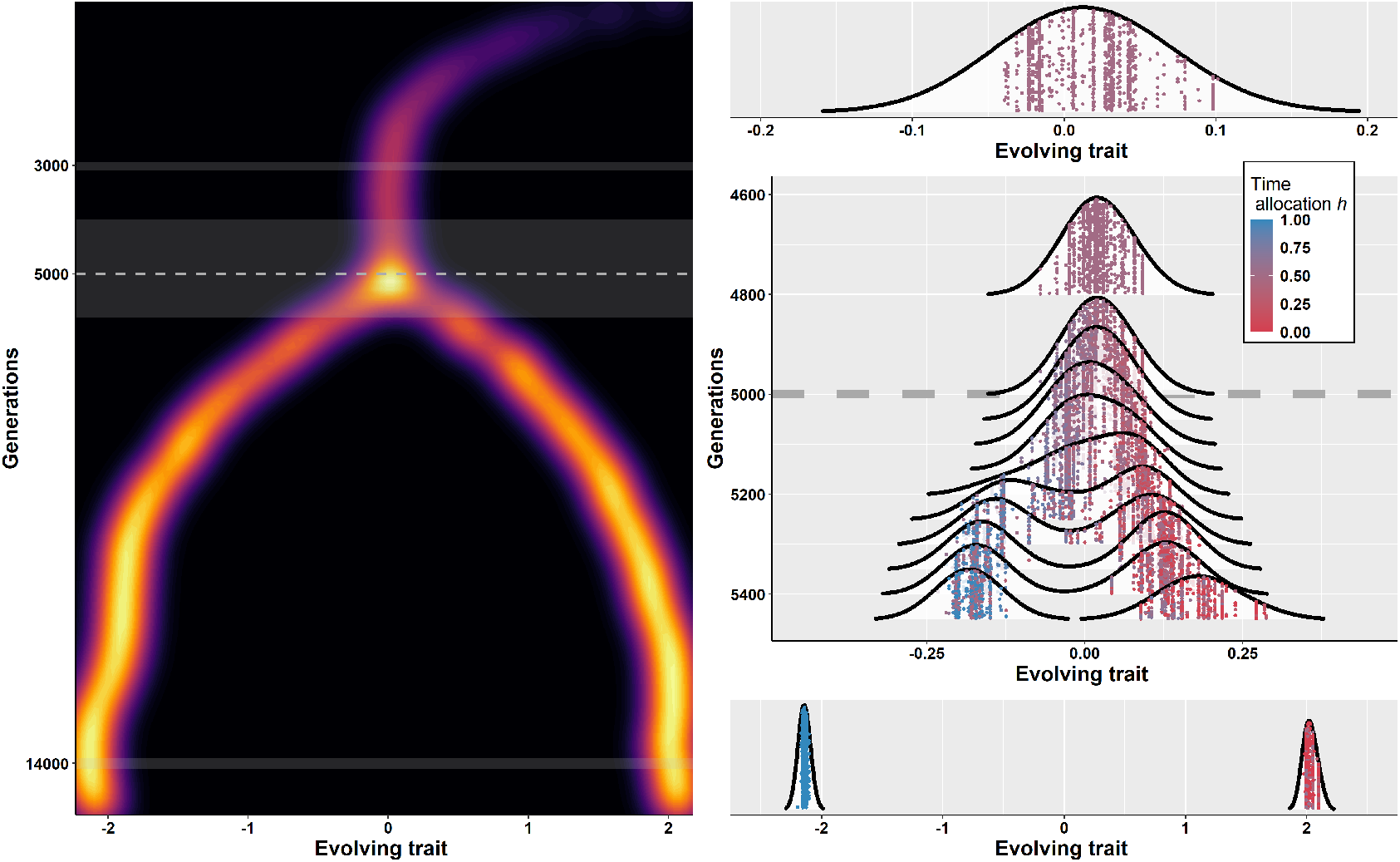
Evolution of genetic diversity and economic specialisation. Same as Figure S10 except that *η* = 0.5. With low value of *η*, economic specialisation remains limited upon introduction of exchange, but appears as individuals become genetically diverse.

**Figure S12.**
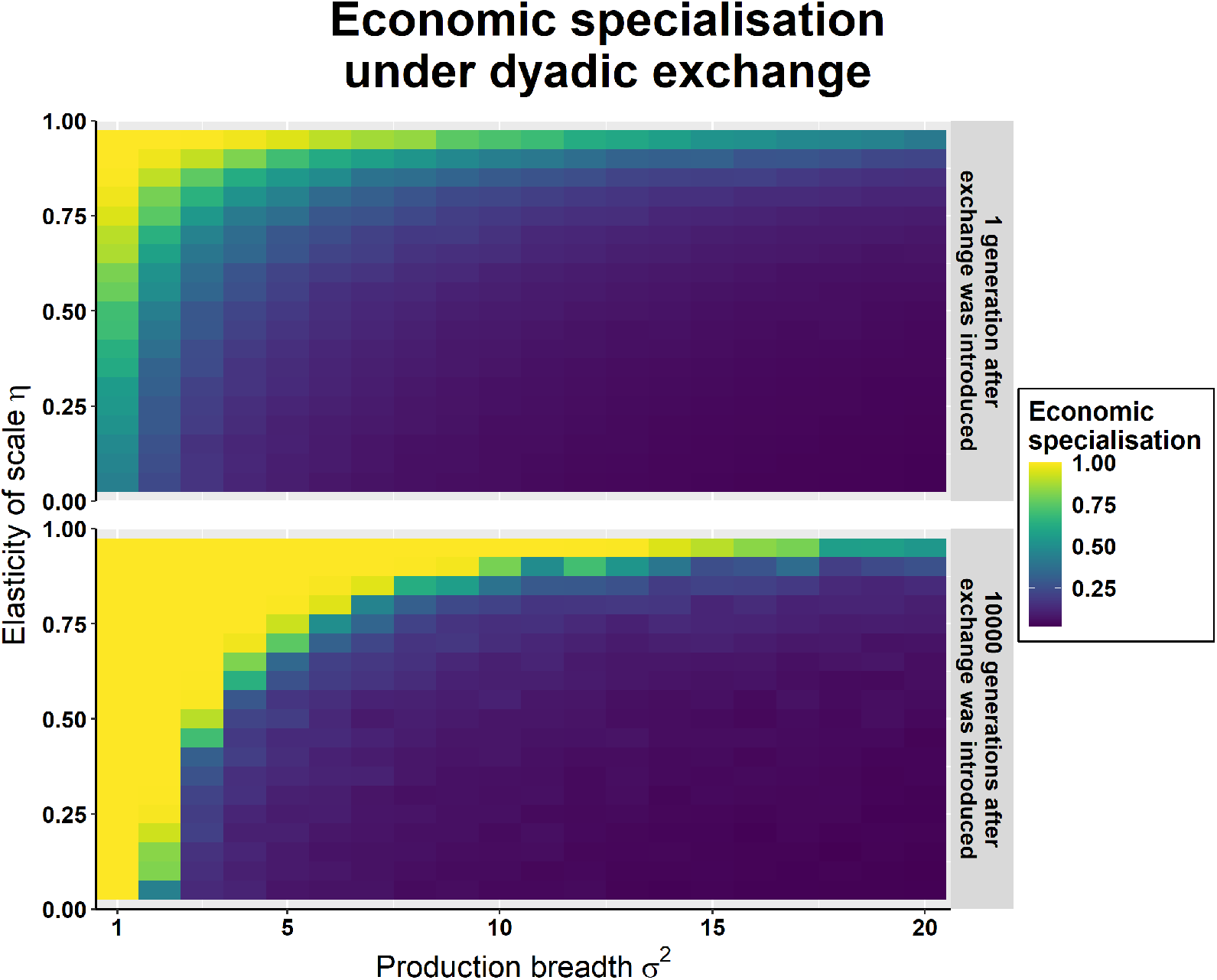
Effect of emergence of exchange on economic specialisation as a function of production breadth *σ*^2^ and elasticity of scale *η*. The panels show average economic specialisation across 10 replicates and after 1 generation (top) and after 10,000 generations (bottom) following the introduction of exchange. Specialisation is measured as the difference between the highest and lowest values of *h* in the population. We consider dyadic exchange where exchange takes place between pairs of isolated individuals. Parameters: equal value of goods *α* = 0.5, optima of production *o*_x_ = −2 and *o*_y_ = 2, population size *N* = 1000, mutation rate *µ*_m_ = 0.01, SD of mutations *σ*_m_ = 0.02. Initial population is drawn from a Normal distribution centered on 0 with SD 0.05.

**Figure S13.**
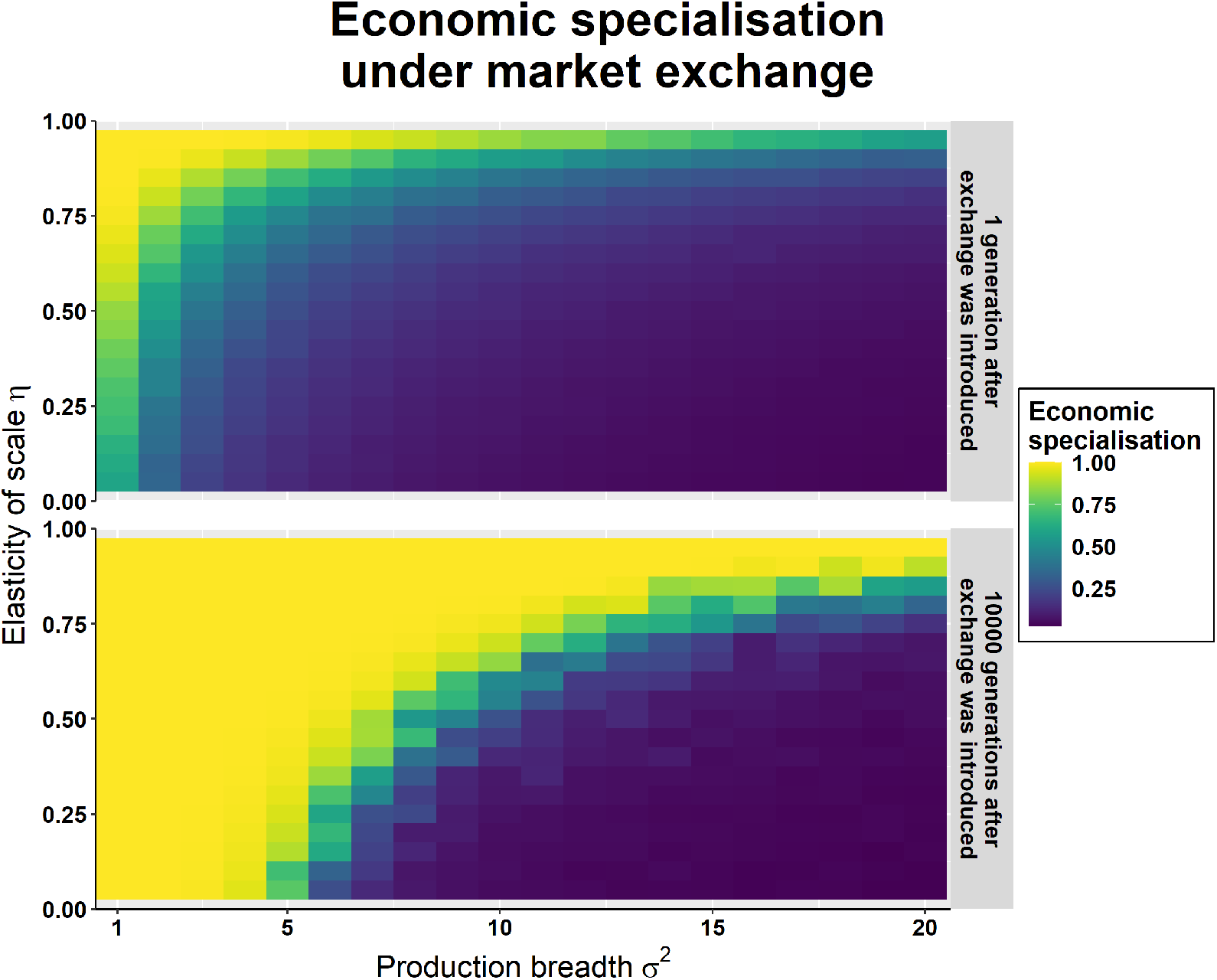
Effect of emergence of exchange on economic specialisation as a function of production breadth *σ*^2^ and elasticity of scale *η*. The panels show average economic specialisation across 10 replicates and after 1 generation (top) and after 10,000 generations (bottom) following the introduction of exchange. Specialisation is measured as the difference between the highest and lowest values of *h* in the population. We consider market exchange where exchange takes place between all individuals. Parameters: equal value of goods *α* = 0.5, optima of production *o*_x_ = −2 and *o*_y_ = 2, population size *N* = 1000, mutation rate *µ*_m_ = 0.01, SD of mutations *σ*_m_ = 0.02. Initial population is drawn from a Normal distribution centered on 0 with SD 0.05.

